# A noncanonical TNL immune hub defines separable recognition and signaling modules for clubroot resistance

**DOI:** 10.1101/2025.10.03.680381

**Authors:** Hao Hu, Fengqun Yu

## Abstract

Plant immunity mediated by nucleotide-binding leucine-rich repeat (NLR) receptors often relies on canonical *EDS1*/*NDR1* signaling, but alternative mechanisms are emerging. We uncover a novel modular, noncanonical immune hub orchestrated by *Rcr1*, a TIR-NLR (TNL) gene conferring clubroot resistance in *Brassica napus* against the root-infecting protist *Plasmodiophora brassicae*. Unlike typical TNLs, Rcr1 engages non-NLR partners in two separable modules: a CP1 (cysteine protease)–WRKY-based recognition module, likely monitoring a pathogen virulence target, and an AP (ankyrin-repeat protein)–ERF-based signaling module, driving jasmonic acid/ethylene-mediated defense. This architecture functions without detectable *EDS1*/*NDR1* involvement, challenging salicylic acid-dominant models of biotrophic immunity and expanding current views of how TNLs can be wired in plant defense. Using high-throughput interactor screening and CRISPR/Cas9 knockouts, we validate these modules, while heat-inducible gene excision reveals *Rcr1*’s critical early role (0–14 days post-inoculation). Together, our findings position *Rcr1* as an exemplar of modular TNL architecture, suggesting that separable recognition and signaling branches may represent a broader principle of immune flexibility in plants. This study redefines TNL flexibility, offering a blueprint for breeding durable disease-resistant crops via modular immune engineering, with clubroot resistance as a model.

## Introduction

Plants have evolved sophisticated immune systems to detect and defend against diverse pathogens. Central to this defense are intracellular nucleotide-binding leucine-rich repeat (NLR) receptors, which perceive pathogen-secreted effectors and initiate robust responses collectively known as effector-triggered immunity (ETI) (Cesari, 2018; Chai et al., 2023). NLRs are broadly divided into two classes based on their N-terminal domains: toll/interleukin-1 receptor (TIR) domain-containing NLRs (TNLs) and coiled-coil-NLRs (CNLs). Canonical signaling mechanisms for these classes have been well characterized in model plants such as *Arabidopsis thaliana*. TNLs typically signal through the Enhanced Disease Susceptibility 1 (EDS1) complex, which interacts with Phytoalexin-Deficient 4 (PAD4) and Senescence-Associated Gene 101 (SAG101) to activate salicylic acid (SA)-dependent defense (Li et al., 2025; Pruitt et al., 2021; Wang et al., 2023a; Xiao et al., 2025). In contrast, CNLs usually operate through Non-race-specific Disease Resistance 1 (NDR1), activating reactive oxygen species (ROS) production, defense gene transcription, and hypersensitive response (HR) (Aarts et al., 1998; Contreras et al., 2023; Day et al., 2006; Knepper et al., 2011; Wang *et al*., 2023a). Recent advances have challenged the exclusivity of these canonical modules, revealing alternative routes of NLR signaling that function independently of EDS1 or NDR1. These noncanonical pathways involve diverse protein partners and signaling nodes, and frequently engage hormonal outputs beyond SA, particularly jasmonic acid (JA) and ethylene (ET). For example, some TNLs have been shown to bypass EDS1 and instead mobilize parallel transcriptional regulators or proteases to activate defense responses (Chai *et al*., 2023). This growing body of work supports a more modular view of NLR function, in which distinct recognition and signaling components can be combined to tailor immune outcomes to specific pathogen lifestyles (Mengiste and Liao, 2025; Sun et al., 2020). As such, the architecture of NLR-mediated resistance is now understood to be more flexible, incorporating both canonical and noncanonical mechanisms with implications for immune specificity, pathway cross-talk, and temporal regulation (Zhang et al., 2017).

Clubroot, caused by the obligate biotrophic protist *Plasmodiophora brassicae*, is one of the most destructive diseases of Brassicaceae crops, posing a major threat to canola (*Brassica napus*) production worldwide. Over 30 clubroot resistance (CR) genes have been mapped or cloned from *Brassica* germplasm, yet few have undergone mechanistic dissection (Chang et al., 2019; Chu et al., 2014; Hatakeyama et al., 2017; Hu et al., 2025; Hu and Yu, 2024; Hu et al., 2023; Huang et al., 2019; Huang et al., 2017; Karim et al., 2020; Li et al., 2024; Rahaman et al., 2025; Ueno et al., 2012; Wei et al., 2024; Yang et al., 2022; Yu et al., 2016; Yu et al., 2017; Yu et al., 2021). Among them, *Rcr1*, a TNL gene originally identified in the *B. rapa* cultivar FN (Chu *et al*., 2014), has been functionally validated and used extensively in Canadian breeding programs via marker-assisted introgression into elite *B. napus* lines (Hu *et al*., 2023). Although *Rcr1*-based resistance has been effective against several *P. brassicae* pathotypes, it has been eroded by the emergence of more virulent strains such as pathotype 5X (Strelkov et al., 2016), underscoring the urgent need to understand the signaling mechanisms that underlie CR durability. Transcriptomic studies have painted a complex picture of the hormonal and transcriptional pathways involved in clubroot resistance. While *Rcr1* in *B. rapa* was shown to activate JA/ET signaling rather than SA (Chu *et al*., 2014), other studies in *B. napus* suggest involvement of *EDS1* and SA-dependent responses (Atem et al., 2024). Additional mechanisms, such as stele-focused defense via the WeiTsing ion channel (Wang et al., 2023b), have further highlighted the diversity of CR strategies across genotypes and tissues. These findings suggest that clubroot resistance may arise from distinct, context-dependent modules with variable reliance on classical immune signaling pathways.

In this study, we investigated the molecular basis of *Rcr1*-mediated resistance to *P. brassicae* in *Brassica napus*, aiming to dissect its signaling architecture and temporal dynamics. By integrating high-throughput interactor screening, CRISPR/Cas9-mediated functional validation, and an inducible gene-excision system, we identified distinct molecular modules operating downstream of *Rcr1*. Our results reveal a noncanonical immune signaling mechanism that bypasses EDS1/NDR1 and instead engages alternative partners to activate JA/ET-mediated defense. These findings support a modular view of TNL function and contribute to the growing paradigm shift in how NLR-mediated resistance is conceptualized in plant-microbe interactions.

## Materials and Methods

### Plant Materials and Growth Conditions

The clubroot-susceptible, spring-type doubled haploid (DH) canola line *Brassica napus* DH12075, originally developed by Dr. Séguin-Swartz (Agriculture and Agri-Food Canada, Saskatoon), was used as the recipient genotype for *Agrobacterium*-mediated transformation of *Rcr1* and subsequent gene editing. Root tissues of pak choi cultivar FN (*B. rapa* ssp. *Chinensis*), the original clubroot-resistant source from which *Rcr1* was cloned, were used to extract total RNA and construct a cDNA library after inoculation with *P. brassicae* pathotype 3H.

All plants were cultivated in a greenhouse under long-day conditions (LDC, 16 h light/8 h dark at 22°C with 230 μmol m⁻² s⁻¹ light at canopy level). Heat treatment (HT) sessions were conducted at 37°C for 30 hours per treatment.

### NGS-Y2H Screening for Rcr1-Interacting Proteins

Protein–protein interactions were identified using a next-generation sequencing–coupled yeast two-hybrid (NGS-Y2H) system with protocol adaptations for high-throughput screening (Fig. 1A) (Suter et al., 2013; Suter et al., 2015).

**Figure 1.**
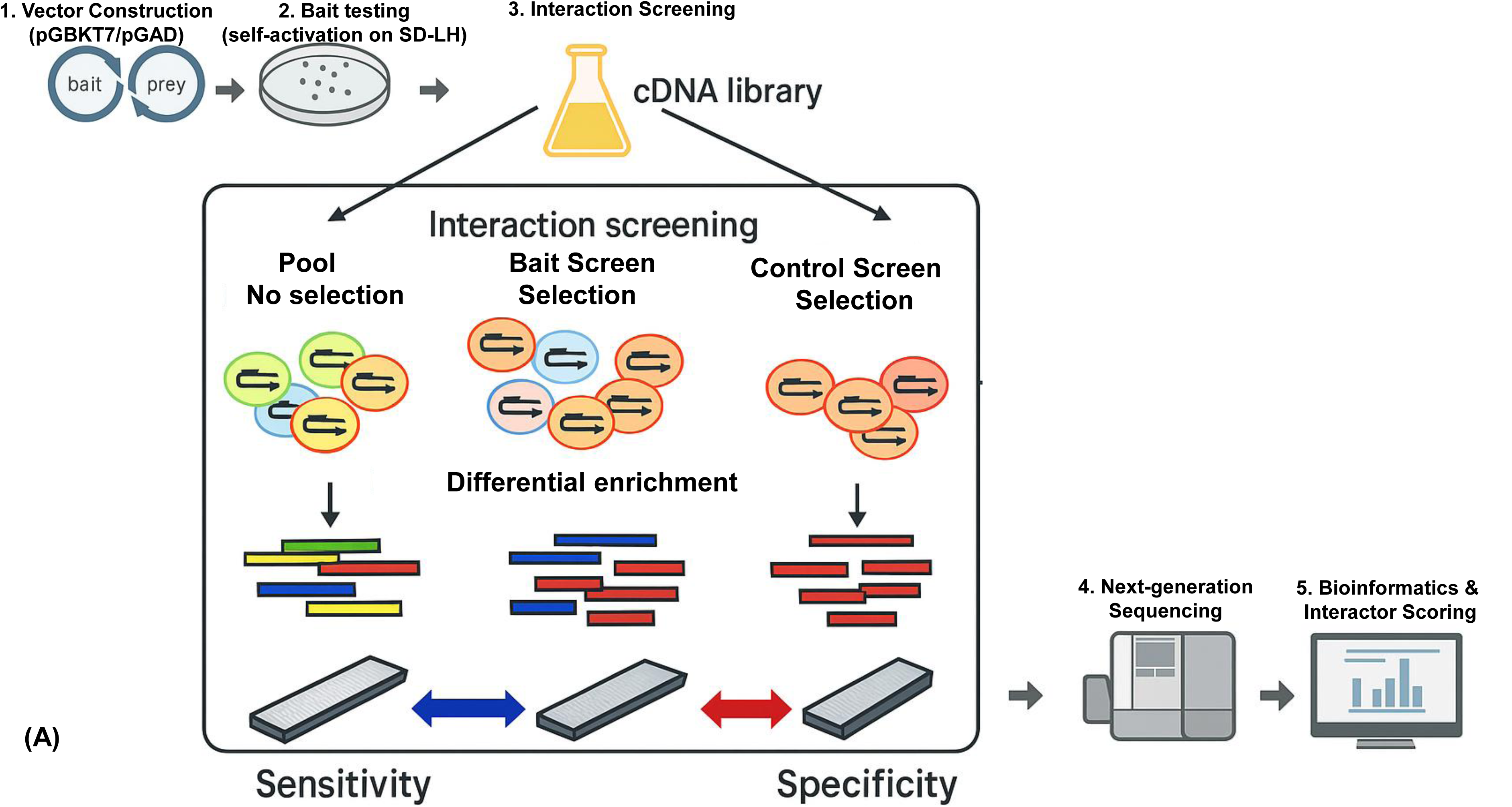
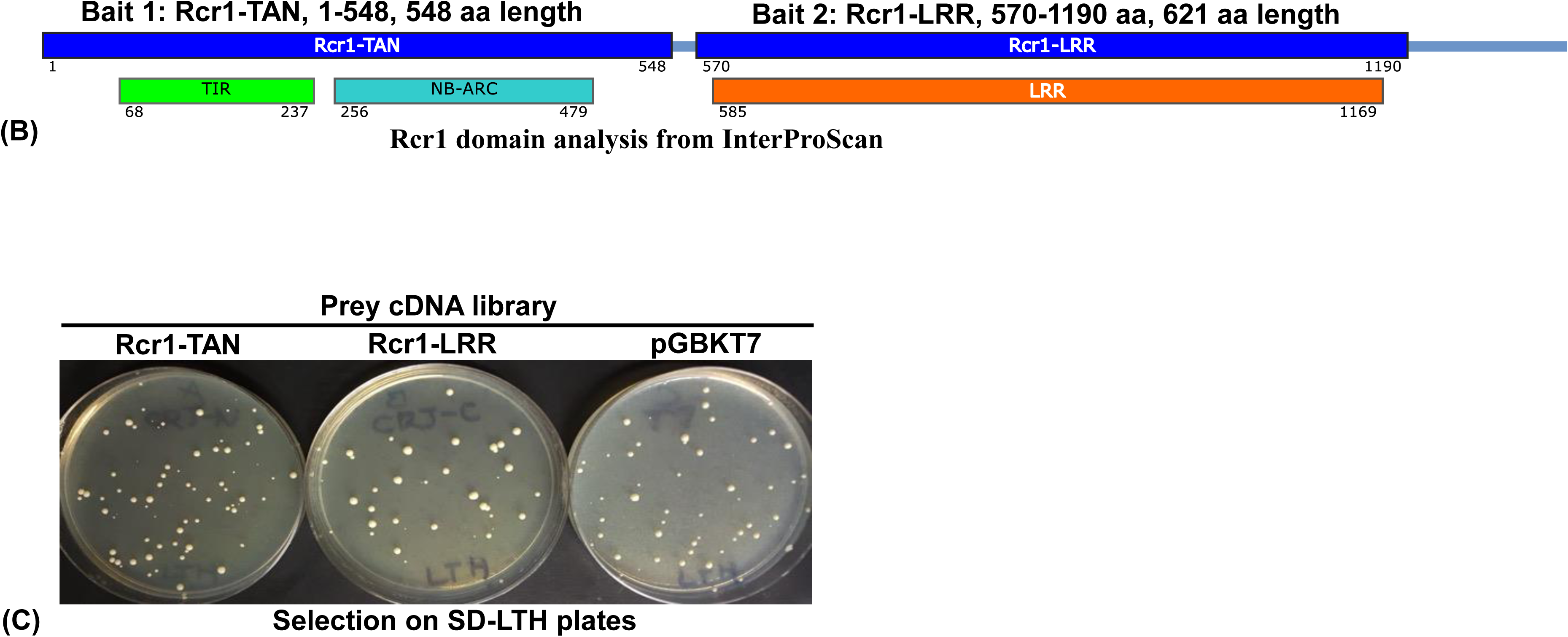
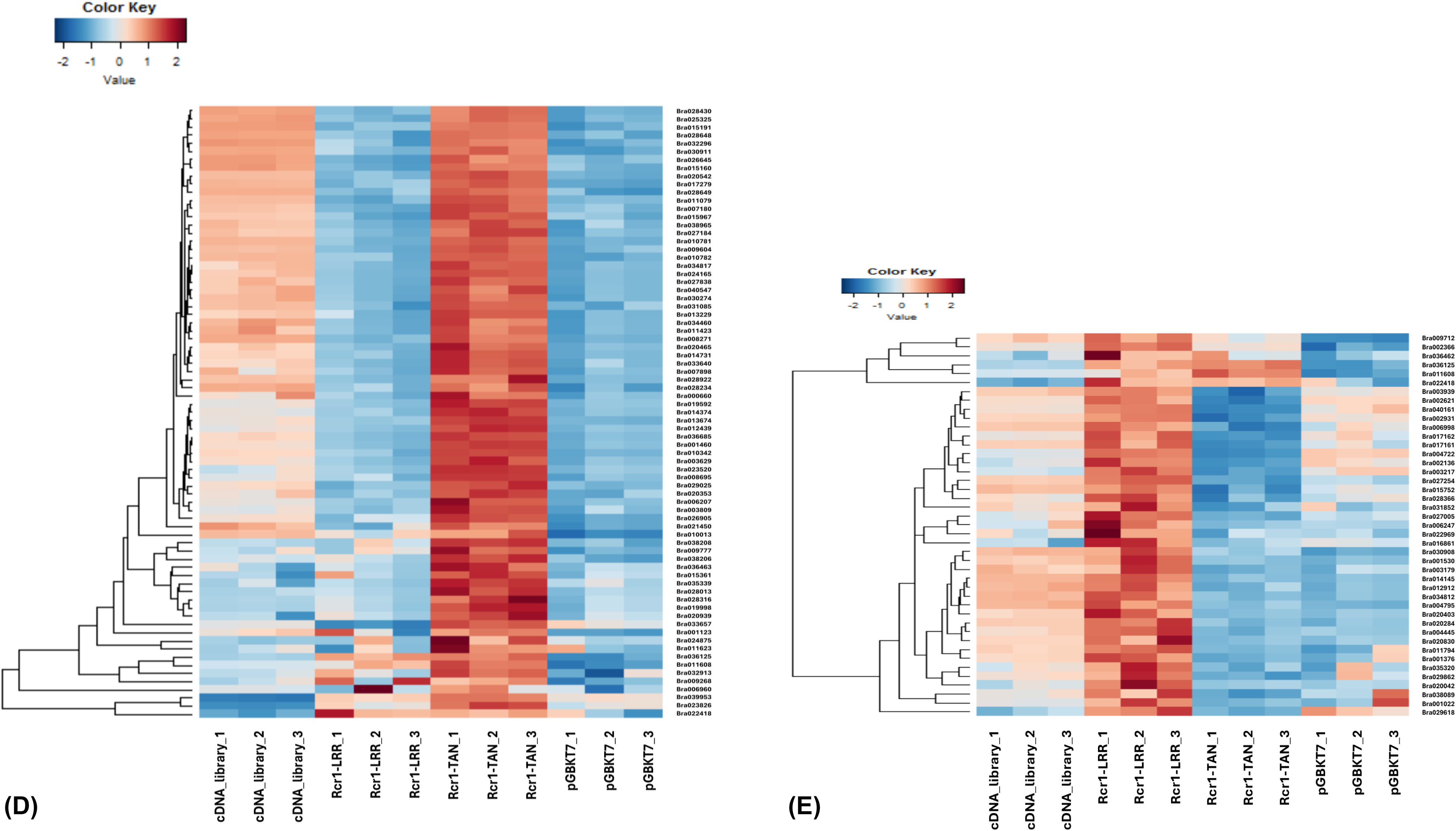
Rcr1-interacting proteins identified via NGS-Y2H. (A) NGS-Y2H workflow. (B) Bait design: Domain analysis (InterProScan) showed Rcr1 as a typical TNL type protein with three core domains of TIR, NB-ARC, and LRR. Two partial constructs (Rcr1-TAN, and Rcr1-LRR) were designed to capture domain-specific interactions. (C) Interaction screening with selection on SD/-Leu/-Trp/-His (SD-LTH) plates. (D) 75 significantly enriched genes over non-selected cDNA library/empty vector control and fulfilling both criteria (log₂FC ≥ 1; FDR ≤ 0.05) were identified as final hits for bait Rcr1-TAN. (E) 43 genes significantly enriched genes over non-selected cDNA library/empty vector control and fulfilling both criteria (log₂FC ≥ 1; FDR ≤ 0.05) were identified as final hits for bait Rcr1-LRR.

Domain annotation of the 1325-aa TNL protein Rcr1 (InterProScan) revealed three core domains: TIR (68–237 aa), NB-ARC (256–479 aa), and LRR (585–1169 aa). To assess domain-specific interactions, two partial constructs, i.e., Rcr1-TAN (1–548 aa, encompassing <u>T</u>IR <u>a</u>nd <u>N</u>B-ARC), and Rcr1-LRR (570–1190 aa, covering LRR), were synthesized (IDT) and cloned into vector pGBKT7 (Takara Bio) via Gibson assembly (NEB) (Fig. 1B). Constructs expressed the bait protein fused to the GAL4 DNA-binding domain (DBD) and carried *TRP1* for selection and a c-Myc tag for validation. The empty pGBKT7 vector served as a control.

The normalized prey library was generated using total RNA extracted from infected roots of *B. rapa* cv. FN collected at 7 days post-inoculation (dpi) with pathotype 3H. Prey cDNAs were cloned into vector pGAD (Takara Bio), which expresses the GAL4 activation domain fusion and carries *LEU2* and an HA tag.

Self-activation was assessed on SD/-Trp/-His (SD-TH) medium. Bait constructs showed negligible activation (<0.0001%), allowing screening without 3-Amino-1,2,4-triazole. Mating of bait and prey yeast strains was conducted in YPDA, and diploids were selected on SD/-Leu/-Trp (SD-LT) medium before being screened for interaction-dependent HIS3 activation on SD/-Leu/-Trp/-His (SD-LTH) (Fig. 1C; Table S1). Triplicate screens were conducted for each bait construct and the vector control (nine screens in total), and the unselected cDNA library was also sequenced in triplicate, bringing the total number of samples for NGS analysis to 12 (Table S2). Surviving colonies were pooled after 5–7 days for DNA extraction. Prey cDNAs were PCR-amplified from yeast genomic DNA, barcoded, and sequenced on an Illumina NextSeq500 (75-bp single-end reads).

Reads were quality-checked (FastQC) and mapped to the *B. rapa* Chiifu v1.5 reference genome (BRAD) using Subjunc (Subread v2.0.3) (Liao et al., 2013a). Gene-level counts were assigned using FeatureCounts (Liao et al., 2013b) and filtered to retain genes with counts per million (CPM) ≥1 in at least two samples. FPKM (fragments per kilobase of transcript per million mapped reads) values were computed for expression analysis.

To assess interaction specificity, gene enrichment was evaluated relative to both the non-selected cDNA library and vector control using the *voom* transformation with linear modeling (*limma*) (Law et al., 2014). Interactors were retained based on log₂ fold-change (log_2_FC) ≥1 and false discovery rate (FDR) ≤ 0.05 (Fig. 1D & 1E). Gene ontology (GO) enrichment was analyzed using AgriGO with *A. thaliana* as the reference, to infer functional roles of high-confidence interactors. Full results are summarized in Supplementary Tables S3 and S4.

### Vector Construction for Plant Transformation

The *Rcr1* expression vector with inducible auto-excision feature, pHHCGR-*Hsp18.2*:*Rcr1*, was developed based on our CRISPR/Cas9 platform for cisgenic breeding (Hu and Yu, 2022) (Fig. S1A). The *Rcr1* coding sequence (4,609 bp) was cloned between the maize *Ubiquitin* promoter (*Pubi*) and *NOS* terminator (*Tnos*), replacing *eGFP* in pHHCGR-*Hsp18.2*. *HygR* replaced the *Bar* selection marker to enable sequential transformation. Heat-inducible excision of *Rcr1* was achieved using *Cas9p* under the control of *Hsp18.2* promoter (Takahashi et al., 1992).

To generate gene knockout (KO) mutants, a dual-sgRNA CRISPR/Cas9 vector (Construct #1) was used (Zhao et al., 2016) (Fig. S1B). Target sgRNA pairs were designed for both homologous alleles in the *B. napus* DH12075 genome using CRISPR-GE (Xie et al., 2017) (Table 1). Deletion sizes were 167 bp for *CP1*, 324 bp for *CP2*, and 249 bp for *AP*. Primers used in this study are listed in Table S5.

**Table 1.**
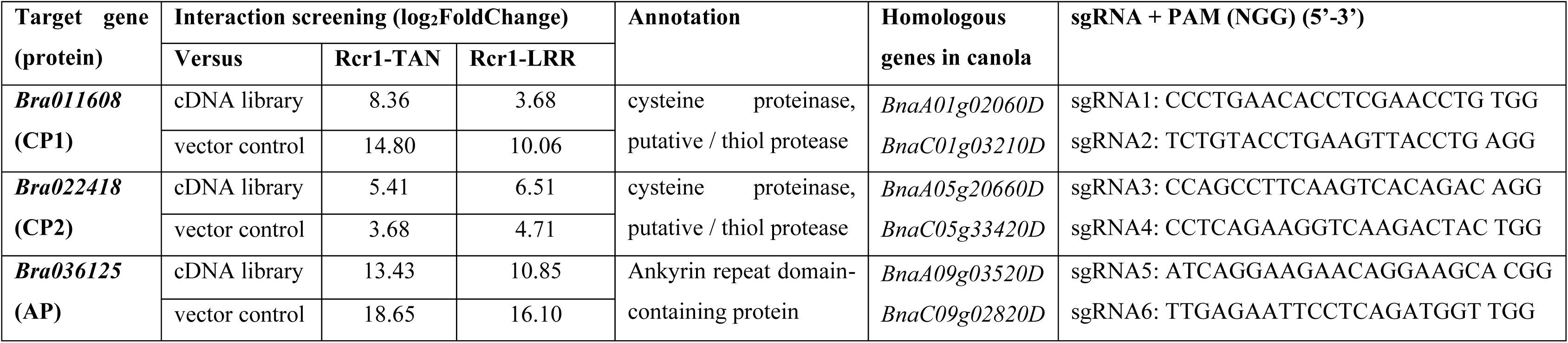
Selection of Rcr1-interacting protein candidates.

### *Agrobacterium*-Mediated Transformation

Vectors were introduced into *Agrobacterium tumefaciens* GV3101 by electroporation. DH12075 was first transformed with pHHCGR-*Hsp18.2*:*Rcr1* to generate homozygous transgenic lines (DH12075+*Rcr1*), selected via ddPCR (Hu *et al*., 2023). One of these lines were used as a recipient for Construct #1. T₀ transformants with biallelic/homozygous mutations were confirmed by PCR and Sanger sequencing, and three independent transformants from each transformation were selected as three biological replicates. Due to the natural limitation of clubroot pathogenicity tests (Hu *et al*., 2023), only T₁ progeny of those selected transformants were used in all functional assays.

### Genotype Panel Establishment

A genotype panel was established with 6 knockout mutants derived from DH12075 and DH12075+*Rcr1*, targeting *CP1*, *CP2*, and *AP*, respectively. Functional status is summarized in Table 2. All genotypes were confirmed by both PCR and sequencing (Fig. S2 & S3). Off-target risks using CRISPR/Cas9-based gene editing were minimized using dual-sgRNA designs and further checked with sequencing (Fig. S4).

**Table 2.**
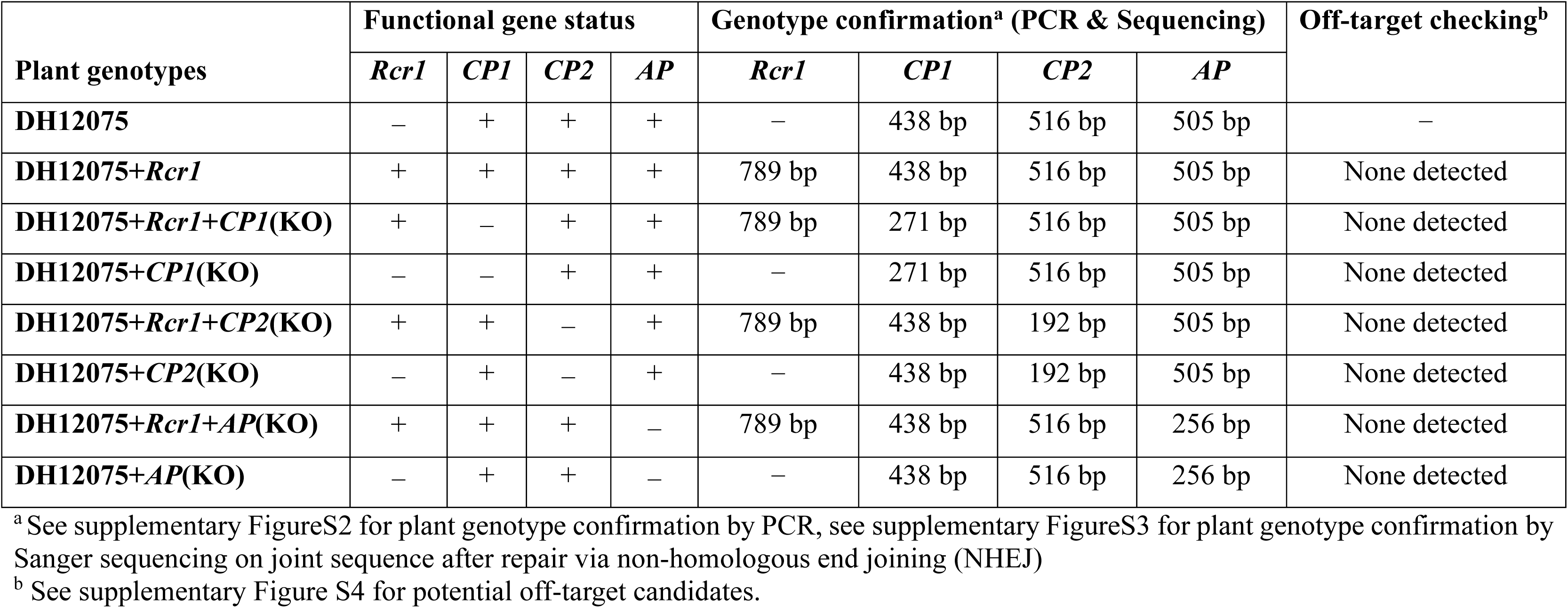
The plant genotype panel used in this study.

### Pathogenicity Tests and Heat Treatments

Pathotype 3H of *P. brassicae*, provided by Dr. Strelkov (University of Alberta), was used for all inoculations. T₁ plants were evaluated using a 0–3 disease rating scale (Kuginuki et al., 1999), and disease severity index (DSI) was calculated (Strelkov et al., 2006). DH12075 and DH12075+*Rcr1* served as susceptible and resistant controls. Heat-induced *Rcr1* excision was applied at 7, 14, or 28 dpi (HT1–HT3), each with 30 hours at 37°C. Six sampling points (S1–S6) were set immediately before and three days after each HT. Disease ratings were conducted at 6 weeks post inoculation (wpi) with three biological replicates per treatment (Fig. 2A). All pathogenicity tests were repeated three times with similar results (Fig. 2B, 2C).

**Figure 2.**
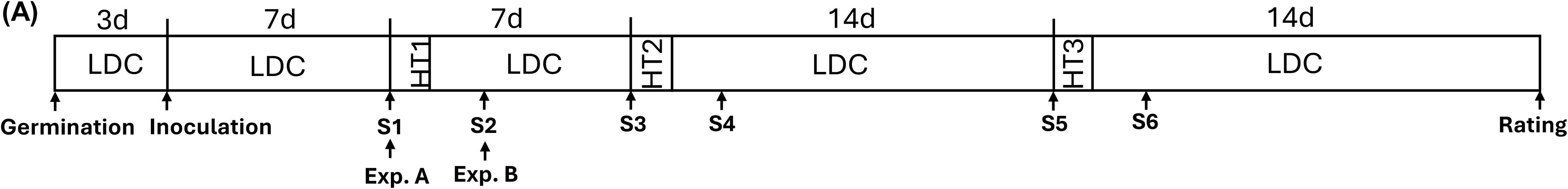
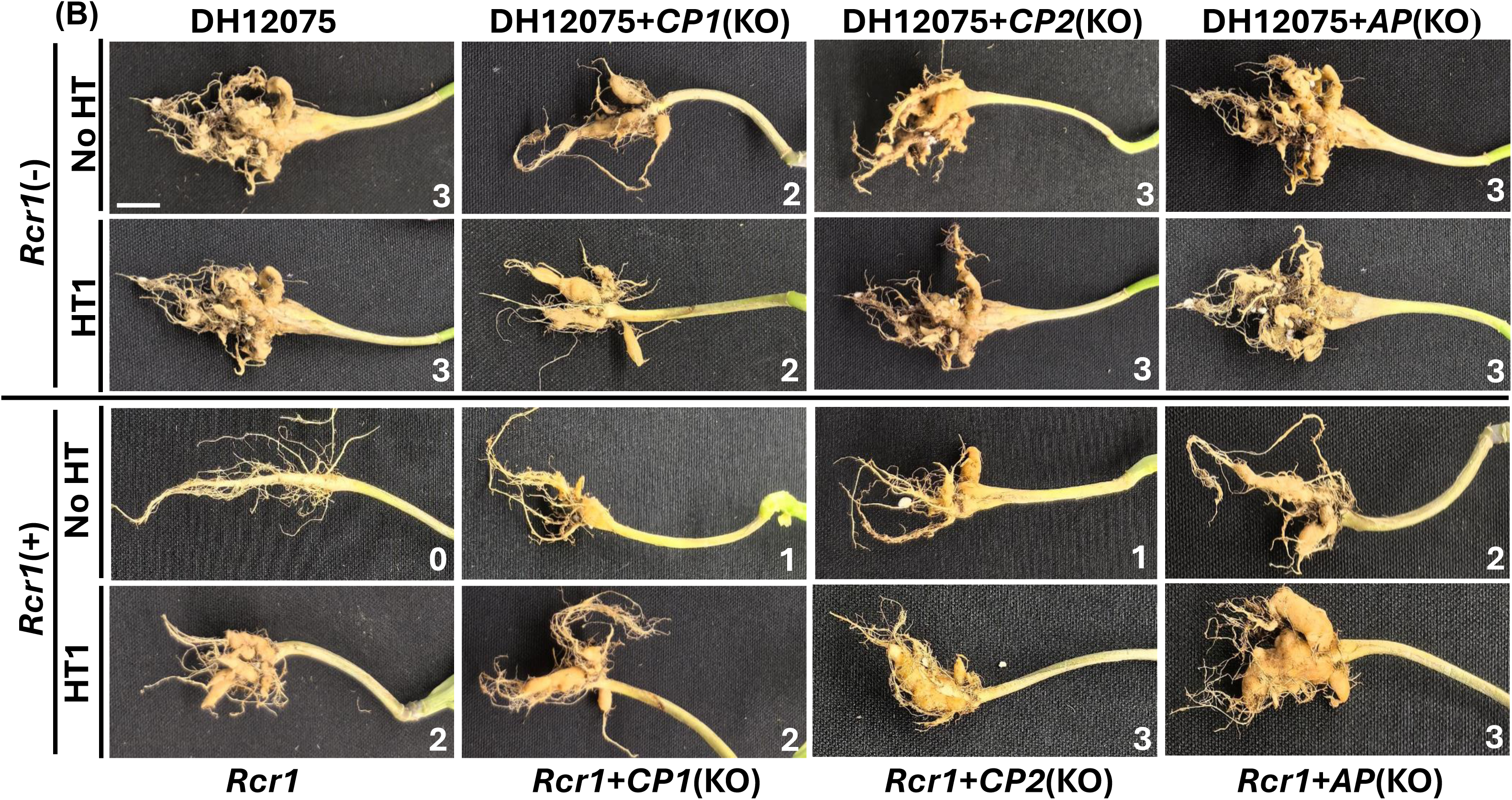
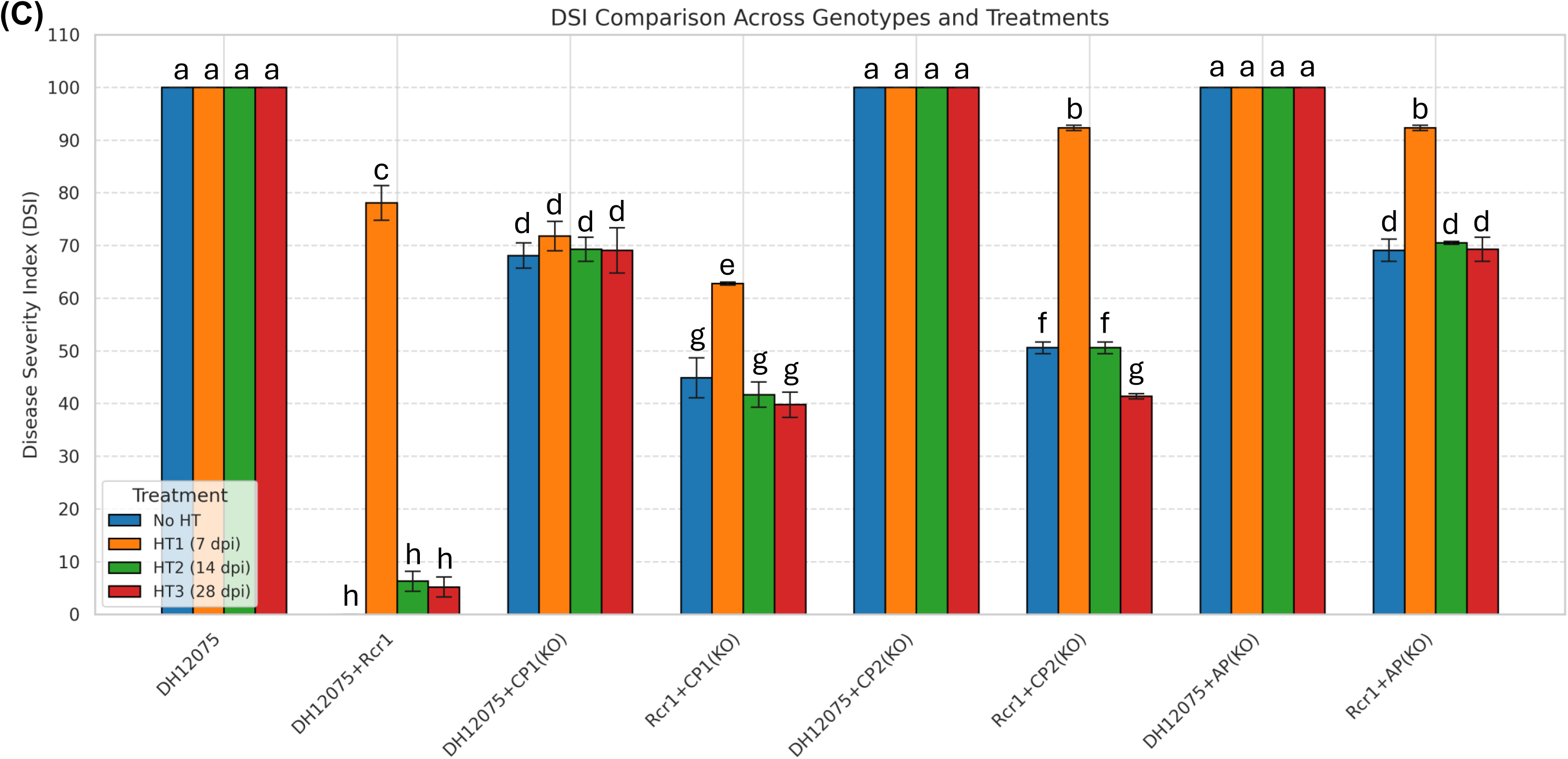
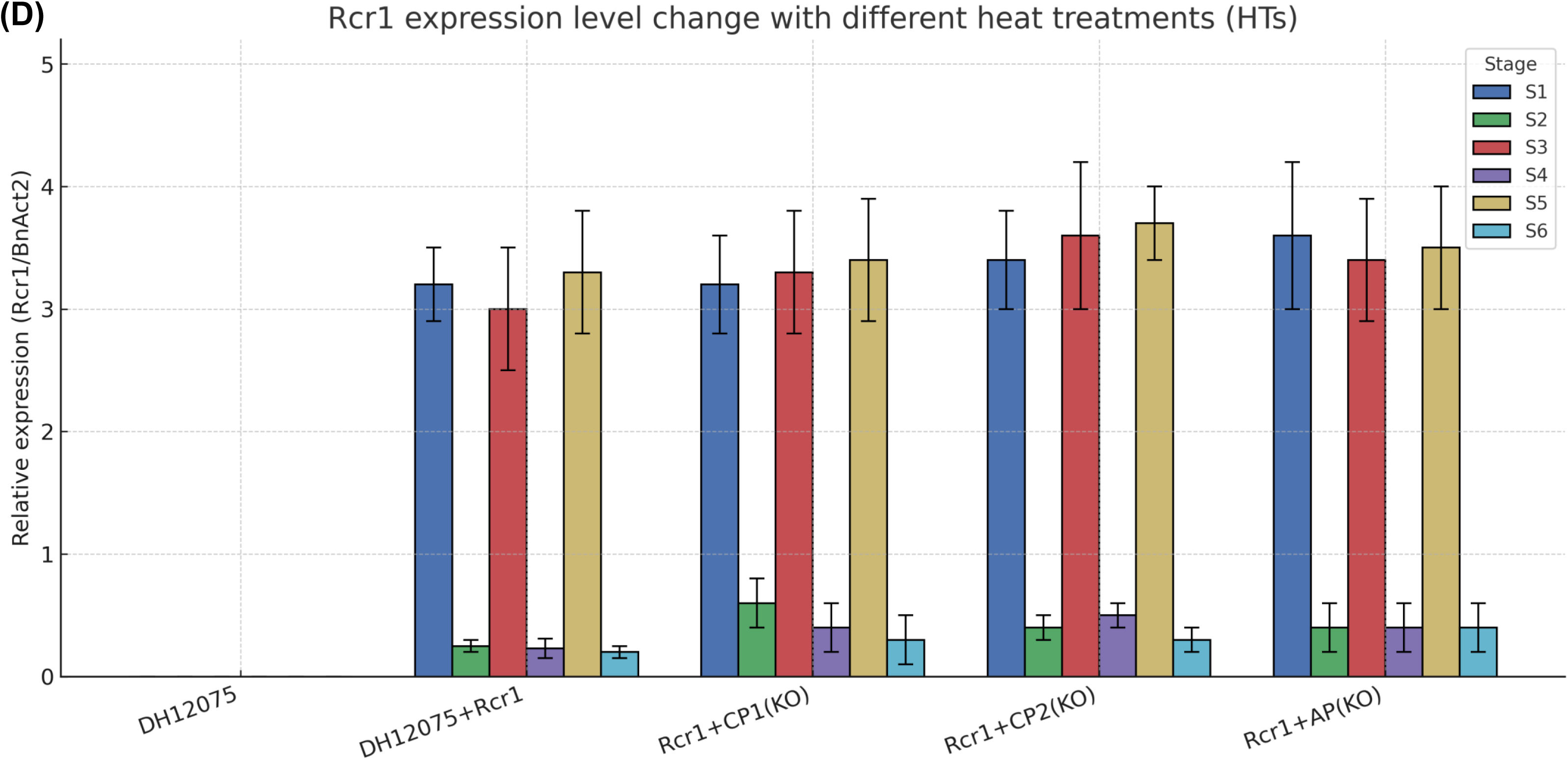
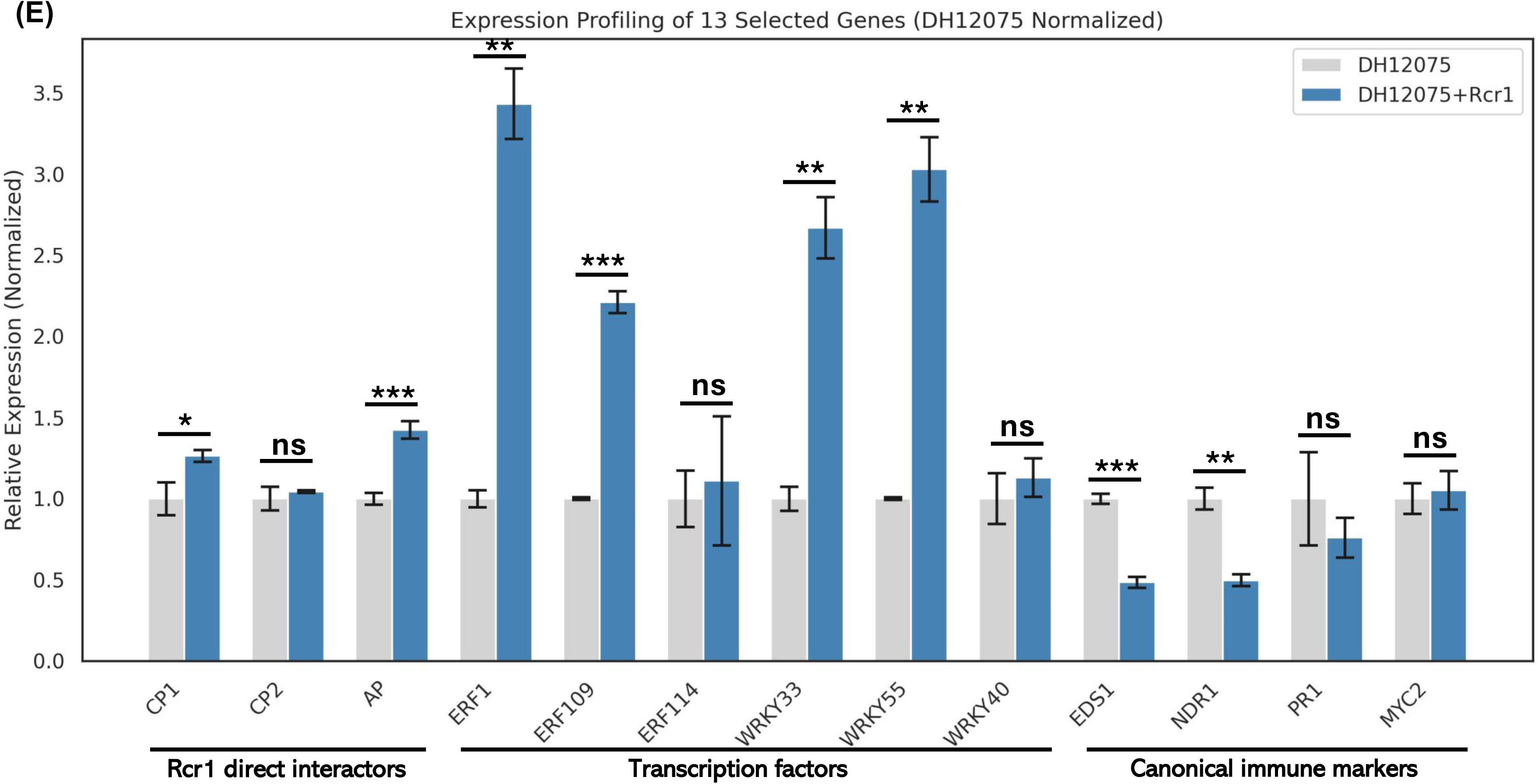
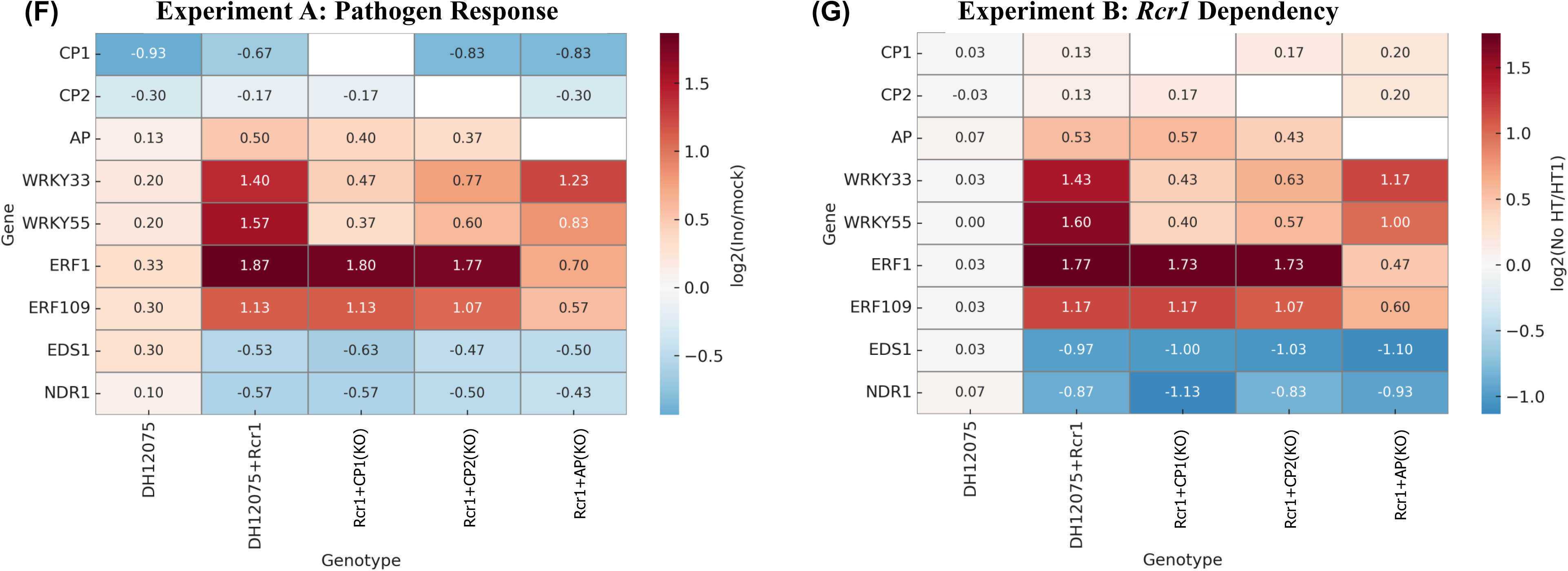
Functional dissection of direct and indirect interactors in *Rcr1*-mediated resistance network. (A) Heat treatments (HTs) were applied to test critical timing for *Rcr1* function and *Rcr1* dependency. LDC, long day condition (16 h light/8 h dark at 22°C with 230 μmol m⁻² s⁻¹ light at canopy level); HT1-HT3, heat treatment 1 to 3; S1-S6, sampling point 1 to 6; d, day. (B) Inoculation results on different genotypes. One representative plant is shown for each genotype-treatment combination, all exhibiting the dominant disease rating (rating scores of 0-3 are labelled on the bottom right corner). Minor phenotypic variation was observed and is detailed in Figure S5. Scale bar, 1 cm. (C) Disease severity index (DSI) comparison across genotypes and treatments. The error bars represent the mean ± standard deviation (SD) (three biological replicates, each replicate with 8-10 plants). Statistical significance (letter a to h) was determined using one-way ANOVA followed by Tukey’s HSD test (*p* < 0.05). (D) *Rcr1* expression level change with different heat treatments. The relative gene expression levels was assessed by RT-qPCR using *BnAct2* as internal reference. Mean and SD were calculated and are presented as bar height and error bars, respectively. (E) Expression profiling of selected genes in DH12075 (*Rcr1*-) and DH12075+*Rcr1* (*Rcr1*+) upon clubroot infection. Statistical analysis was conducted using Student’s *t*-test based on three independent biological replicates for each genotype. Mean and SD were calculated and are presented as bar height and error bars, respectively. Statistical significance was assigned using the following thresholds: ns, not significant; *, *p* < 0.05; **, *p* < 0.01; ***, *p* < 0.001. (F) Experiment A: Pathogen response of *Rcr1*-related key genes monitored via RT-qPCR. (G) Experiment B: *Rcr1* dependency of *Rcr1*-related key genes revealed via RT-qPCR.

### RT-qPCR Analysis

The relative gene expression levels were assessed by RT-qPCR using *BnAct2* as the internal reference gene. For *Rcr1* expression analysis, pooled new leaves were collected at defined sampling points (S1–S6) and used for total RNA extraction with the RNeasy Plant Mini Kit (Qiagen) (Fig. 2D). To conduct gene expression profiling in *Rcr1*(+) and *Rcr1*(–) backgrounds (i.e., DH12075+*Rcr1* vs. DH12075) at S1 (Fig. 2E), and to further examine expression changes of nine *Rcr1*-associated genes (*CP1*, *CP2*, *AP*, *WRKY33*, *WRKY55*, *ERF1*, *ERF109*, *EDS1*, and *NDR1*) (Fig. 2F & 2G), total RNA was extracted from pooled root tissues of each biological replicate (8– 10 plants per replicate). For Experiments A and B, samples were collected at S1 (inoculated vs. mock) and S2 (no HT vs. HT1), respectively (Fig. 2A). Relative expression levels were calculated using two approaches depending on the experimental design: the 2^–ΔCt method was applied for *Rcr1* expression dynamics and general gene profiling; the 2^–ΔΔCt method was used to quantify pathogen-induced changes and *Rcr1*-dependent expression responses. RT-qPCR protocols followed Hu et al. (2023) and primer sequences are listed in Table S5.

### PCR and Sanger Sequencing

PCR was used to validate transgene presence, gene deletions, and absence of off-target edits (Fig. S2). Genomic DNA was extracted from new leaves of individual plant using the DNeasy Plant Mini Kit (Qiagen). Potential off-target sites were predicted using CRISPR-GE. Amplicons were cloned into pCR2.1 (Invitrogen) and sequenced (NRC, Saskatoon). PCR protocols followed Hu et al. (2023) and primer are listed in Table S5.

### Statistical Analysis

Statistical analyses on gene expression profiling experiment (Fig. 2E) and pathogenicity test (Fig. 2C) were performed using Student’s *t*-test and one-way ANOVA followed by Tukey’s HSD test (*p* < 0.05), respectively. All statistical analyses were performed using R (version 4.3.1).

## Results

### NGS-Y2H screening identifies high-confidence Rcr1-interacting proteins

To elucidate the molecular components involved in *Rcr1*-mediated clubroot resistance, we performed high-throughput NGS-Y2H screening (Fig. 1A) using the TIR-and-NBS (Rcr1-TAN) and LRR (Rcr1-LRR) domains of Rcr1 as baits (Fig. 1B). Interaction screens with both baits and an empty vector control revealed comparable background activation levels (∼0.01%), validating the stringency of the assay and the suitability of our system for identifying specific interactors (Fig. 1C, Table S1).

From the Rcr1-TAN screens, 87 genes were significantly enriched over the non-selected cDNA library and 480 over the vector control, with 75 genes fulfilling both criteria (log₂FC ≥ 1; FDR ≤ 0.05) (Fig. 1D). The Rcr1-LRR screens yielded 147 genes enriched against the cDNA library and 239 against the vector control, with 43 overlapping hits (Fig. 1E). These candidates were filtered to exclude vector-specific and non-reproducible signals, resulting in 75 and 43 high-confidence interactors from the Rcr1-TAN and Rcr1-LRR baits, respectively (Tables S3 and S4).

Among these, three interactors, CP1 (papain-like cysteine protease, PLCP), CP2 (a homologous PLCP), and AP (ankyrin-repeat protein), were detected with both baits across independent screens and satisfied the enrichment thresholds with high reproducibility (Table 1). Functional annotation via GO analysis suggested functions of protease activity (CP1 and CP2) and signal transduction (AP), consistent with potential roles of effector-monitoring (guardee) or signaling mediator, respectively (Table S3 and S4). These three proteins were prioritized for further validation based on their interaction specificity, expression during infection, and relevance to immune signaling pathways (Table 1). Since functional validation was performed in canola, the homologous *B. napus* genes corresponding to the three prioritized Rcr1-interacting proteins were used for sgRNA design to generate knockout mutants (Genoscope v5, CRISPR-GE).

### Generation of a genotype panel for functional dissection of *Rcr1* pathway

To validate the functional relevance of Rcr1 and its interactors, we established a genotype panel consisting of eight *B. napus* lines, including wild-type DH12075, *Rcr1*-transformed DH12075 (DH12075+*Rcr1*), and six CRISPR/Cas9-derived knockout lines targeting *CP1*, *CP2*, and *AP* in both the *Rcr1*(+) and *Rcr1*(–) backgrounds (Table 2). All knockouts were generated using dual-sgRNA constructs designed to delete coding sequences of target genes (167-324 bp), and confirmed via PCR and Sanger sequencing (Fig. S2, S3). Off-target effects were minimized using *in silico* prediction (CRISPR-GE) and were not detected in edited lines (Fig. S4).

All genotypes in the panel were homozygous at the target loci and displayed reproducible phenotypic trends across independent biological replicates, with statistically significant differences consistently observed between key comparisons (Fig. 2B). The panel enabled systematic dissection of gene-specific contributions under both *Rcr1*(+) and *Rcr1*(–) conditions, despite natural biological variability inherent to clubroot pathosystems (Fig. S5).

### Functional roles of Rcr1-interacting proteins in clubroot resistance

To examine the contribution of each gene to disease resistance, all eight genotypes were subjected to *P. brassicae* pathotype 3H inoculation and evaluated at 6 wpi. Heat treatments were applied at 7, 14, or 28 dpi, corresponding to early, mid, and late infection stages (HT1–HT3), and sampling points were set before and after each HT (S1–S6) (Fig. 2A).

In the absence of *Rcr1* (DH12075), plants showed severe disease symptoms with a DSI of 100% (Fig. 2B). In contrast, DH12075+*Rcr1* consistently displayed full resistance (DSI = 0%), validating the protective effect of the transgene. The *Rcr1*-transgenic line was created using our self-excising CRISPR vector (pHHCGR-*Hsp18.2*:*Rcr1*, Fig. S1A), enabling heat-inducible removal of *Rcr1* through *Cas9p* expression. Upon heat treatment (37°C for 30 hours) at defined time points, *Rcr1* excision efficiency consistently exceeded 90%, as demonstrated previously (Hu and Yu, 2022) and further validated here by RT-qPCR (Fig. 2D) and loss of genomic presence (Fig. S2). This design allowed temporal resolution of *Rcr1* activity and assessment of downstream gene dependency. Removal of *Rcr1* at 7 dpi (HT1) resulted in partial loss of resistance, whereas excision at 14 dpi (HT2) or later (28 dpi, HT3) retained near-complete resistance. These results demonstrate that *Rcr1* activity is critical during early infection, most likely within the first 0-14 dpi window (Fig. 2C).

In *Rcr1*(+) backgrounds, knockout of *CP1*, *CP2*, or *AP* led to significant increases in DSI compared to DH12075+*Rcr1*, confirming that all three interactors are required for full resistance. However, functional distinctions emerged when analyzing their effects in the *Rcr1*(–) background. *CP1* knockout in DH12075 resulted in a modest yet significant reduction in DSI, suggesting that *CP1* is targeted by the pathogen independently of *Rcr1* and may act as a direct effector target (i.e., a guardee). In contrast, knockout of *CP2* or *AP* in the absence of *Rcr1* had no effect on disease progression, indicating their functions are *Rcr1*-dependent (Fig. 2C).

### RT-qPCR profiling reveals dual-modular noncanonical defense activation

To delineate the downstream signaling architecture of *Rcr1*-mediated resistance, we conducted expression profiling of immunity-related genes including marker genes of JA/ET/SA-pathways. A total of 34 selected genes were analysed via RT-qPCR in both DH12075+*Rcr1* and DH12075 to reveal *Rcr1*-related transcriptional perturbation at early stage after clubroot inoculation (i.e., 7 dpi). As a result, the expression levels of 4 transcription factors (TFs) that usually involved in JA/ET-pathway (Birkenbihl et al., 2012; Cai et al., 2014; Cheng et al., 2013; Kang et al., 2024; Mao et al., 2011; Wang et al., 2020b; Zheng et al., 2006), i.e., *ERF1*, *ERF109*, *WRKY33*, and *WRKY55*, were identified to have significant elevation in DH12075+*Rcr1* comparing to those in DH12075 in response to the inoculation of *P. brassicae* pathotype 3H; however, some canonical marker genes in TNL-mediated plant immunity (Li *et al*., 2025; Wang *et al*., 2023a), e.g., *EDS1*, *PAD4*, *Activated Resistance 1* (*ADR1*), *Isochorismate Synthase 1* (*ICS1*), and *Pathogenesis-Related Protein 1* (*PR1*), were found to be suppressed or unchanged (Fig. 2E, Fig. S6). These RT-qPCR profiling results were supported by the previous RNA-seq study using *Rcr1*-carrying *B. rapa* material (Chu *et al*., 2014). However, several other TFs identified by the RNA-seq profiling were not confirmed by our RT-qPCR in *B. napus* background (Fig. S6), thus they were not included in the following investigations.

The expression pattern of nine key genes was further investigated across inoculated and mock-treated conditions (Fig. 2F, Experiment A: Pathogen response), and with or without *Rcr1* excision (Fig. 2G, Experiment B: *Rcr1* dependency). Genes included the three *Rcr1*-interactors (*CP1*, *CP2*, *AP*), two pairs of TFs (*WRKY33/WRKY55*, *ERF1/ERF109*) linked to recognition and signaling, and two classical resistance gene signaling components (*EDS1*, *NDR1*).

In Experiment A, inoculation of DH12075+*Rcr1* triggered strong induction of *WRKY33*, *WRKY55*, *ERF1*, and *ERF109*, with minimal expression of *EDS1* and *NDR1*, suggesting activation of JA/ET-associated defense rather than canonical TNL/CNL pathways. Expression of *WRKY33* and *WRKY55* was significantly suppressed in *CP1* and *CP2* knockout lines, while *ERF1* and *ERF109* were selectively reduced in the *AP* knockout. These patterns suggest a modular structure wherein *CP1*/*CP2* mediate upstream recognition (via *WRKY* TFs) and *AP* mediates downstream signaling (via *ERF* TFs) (Fig. 2F). Notably, expression of *CP1* and *CP2* was reduced upon pathogen inoculation, while *AP* was not.

In Experiment B, the removal of *Rcr1* at 7 dpi (HT1) substantially reduced the expression of all four JA/ET marker genes (*WRKY33*, *WRKY55*, *ERF1*, *ERF109*) in the DH12075+*Rcr1* background, validating their *Rcr1*-dependency. Out of the three Rcr1-interacting proteins, AP showed significant higher *Rcr1* dependency, consistent with the pathogen response in Experiment A. The selective connection of CP–WRKY or AP–ERF was observed and thus confirmed again. *EDS1* and *NDR1* remained suppressed, but the suppression effect was reduced, further supporting a noncanonical signaling route (Fig. 2G). These findings reveal that *Rcr1* orchestrates a dual-module immune response, comprising a CP1–WRKY*-*based recognition axis and an AP–ERF-based signaling arm, operating independently of *EDS1*/*NDR1* (Fig. 3).

**Figure 3.**
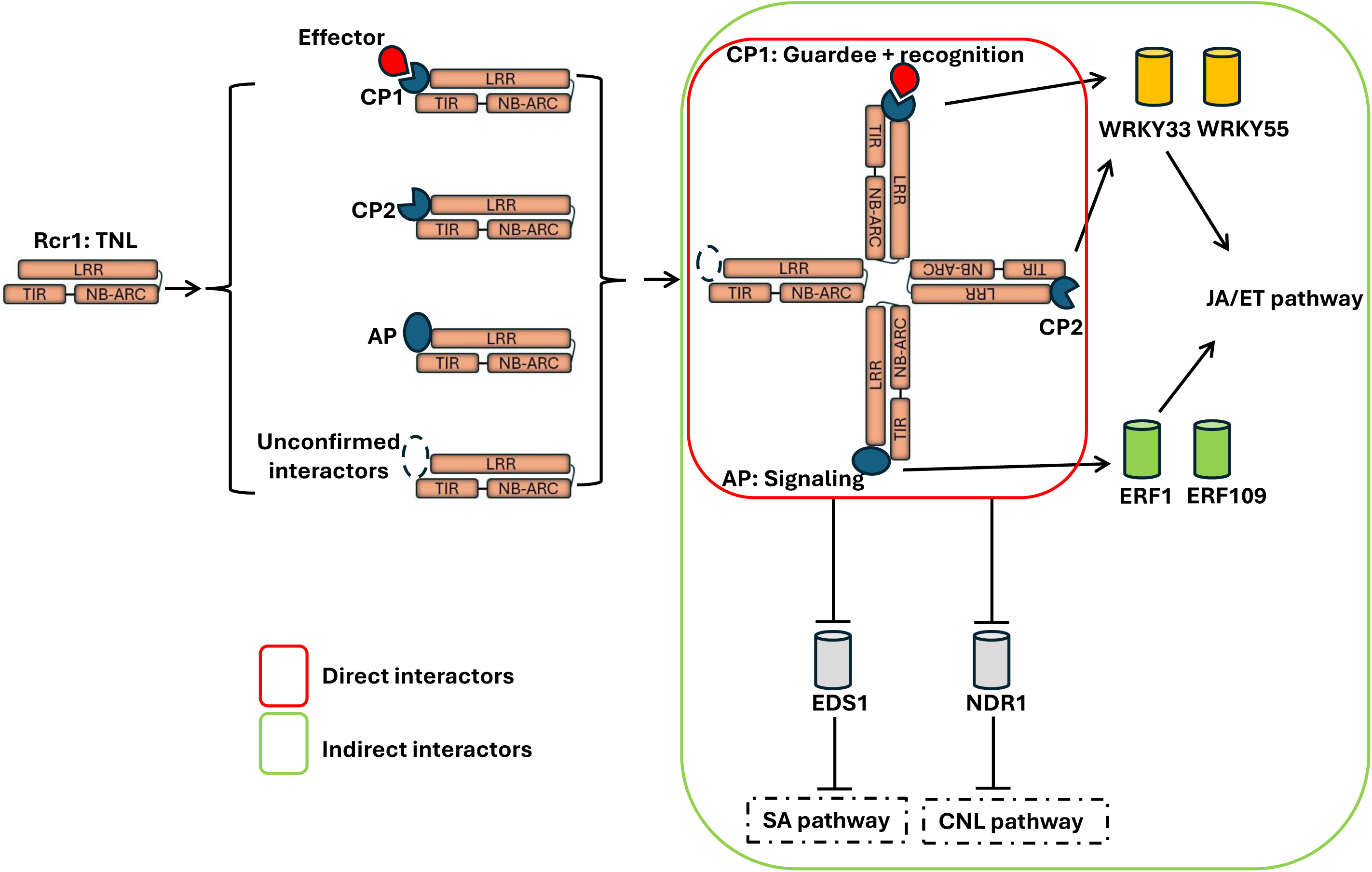
Rcr1 functions as a modular immune hub in a non-canonical TNL network. At early stage post inoculation (i.e., 7 dpi) with *Plasmodiophora brassicae* pathotype 3H, Rcr1 engages three direct protein interactors, i.e., CP1, CP2, and AP, identified via NGS-Y2H screening. Functional assays show that CP1 and AP are required for full resistance, forming two discrete branches of a modular immune network: a CP1–WRKY-based recognition module and an AP– ERF-based signaling module. These branches converge on JA/ET-responsive defense activation. Six downstream proteins, i.e., WRKY33, WRKY55, ERF1, ERF109, EDS1, and NDR1, are considered indirect interactors based on transcriptional response patterns rather than physical association with Rcr1. WRKYs and ERFs exhibit branch-specific expression dependence, while EDS1 and NDR1 are transcriptionally suppressed and appear excluded from Rcr1-mediated signaling. This modular configuration defines a noncanonical TNL signaling route independent of classical EDS1- or NDR1-mediated pathways.

## Discussion

Classical models of plant immunity distinguish defense signaling pathways based on pathogen lifestyle. SA signaling is generally associated with defense against biotrophic and hemibiotrophic pathogens, facilitating localized and systemic acquired resistance (SAR) through hypersensitive cell death and activation of pathogenesis-related (PR) genes. In contrast, JA and ET signaling pathways are typically linked to defense against necrotrophic pathogens and herbivorous insects, supporting the production of antimicrobial compounds and defense-related gene expression without necessarily invoking cell death (Glazebrook, 2005; Pieterse et al., 2009). However, this dichotomy has been challenged by recent studies, particularly in roots, where JA/ET signaling has been shown to play unexpectedly central roles in defense against biotrophs(Mengiste and Liao, 2025; Millet et al., 2010). In clubroot, a root-specific disease caused by the obligate biotroph *P. brassicae*, transcriptomic analyses have revealed the activation of both SA- and JA/ET-associated pathways across different host genotypes and infection stages, indicating that immune signaling is dynamic, tissue-specific, and genotype-dependent (Atem *et al*., 2024; Chu *et al*., 2014; Wang et al., 2022). Within this complexity, our study demonstrates that the TNL gene *Rcr1* mediates resistance in *B. napus* through a distinct pathway that functions without detectable involvement of canonical SA-dependent TNL nodes and instead organizes defense through separable recognition and signaling modules. This highlights that TNLs are not universally dependent on *EDS1*/*NDR1*, expanding current views of how these receptors can be wired.

Our NGS-Y2H screen, employing both the TAN and LRR domains of Rcr1 as baits, identified CP1, CP2, and AP as dual-domain interactors, prioritized based on enrichment strength and biological relevance. Functional validation using CRISPR/Cas9 knockouts showed that *CP1* is essential for *Rcr1*-mediated resistance. Interestingly, its disruption also conferred partial resistance in *Rcr1*(–) backgrounds, suggesting that CP1 may act as a virulence target exploited by *P. brassicae*, as well as a guarded component monitored by Rcr1. While direct interaction between CP1 and pathogen effectors has not yet been demonstrated, previous studies have implicated homologous PLCPs such as AtXCP1 in *Arabidopsis* as effector targets of *P. brassicae* (Pérez-López et al., 2021), supporting CP1’s role as a key recognition node. Additionally, *CP1* was transcriptionally downregulated upon infection, suggesting that it is suppressed by pathogen activity, consistent with its hypothesized role as a guarded virulence target (Pérez-López *et al*., 2021). In contrast, *AP* knockouts did not affect *CP1* or *WRKY* expression, but specifically abolished induction of *ERF1* and *ERF109*, indicating that AP functions downstream of recognition and plays a key role in activating JA/ET signaling. Although the precise biochemical activity of AP remains to be elucidated, its ankyrin-repeat domain is consistent with a role as a scaffolding protein involved in signal transduction (Kolodziej et al., 2021; Wang et al., 2020a). The participation of non-NLR co-factors in TNL signaling has gained attention (Mengiste and Liao, 2025), but remains experimentally underexplored, and our identification of AP represents a timely and significant contribution to this emerging field. CP2 exhibited a minor phenotype, contributing to resistance only in the presence of Rcr1, suggesting a redundant or conditionally relevant role. Notably, disruption of either *CP1* or *AP* did not fully abolish resistance, indicating that additional *Rcr1*-dependent modules exist. Indeed, our interactome analysis identified multiple high-confidence interactors that await functional characterization (Table S3 & S4), reinforcing the concept that Rcr1 operates as a flexible immune hub with a layered or partially redundant architecture.

RT-qPCR profiling further supported this architecture by identifying six indirect interactors that responded to *P. brassicae* infection in an *Rcr1*-dependent manner: *WRKY33*, *WRKY55*, *ERF1*, *ERF109*, *EDS1*, and *NDR1*. These were selected from an initial panel of 34 candidates based on expression specificity, magnitude, and relevance to defense signaling. When tested in the context of *CP1* and *AP* knockouts, *WRKY33* and *WRKY55* were downregulated in *CP1* mutants, whereas *ERF1* and *ERF109* were specifically diminished in *AP* mutants, delineating a CP1–WRKY axis for recognition and an AP–ERF axis for signaling. These transcriptional trends in *B. napus* were consistent with patterns observed in our RNA-seq data from *Rcr1*-carrying *B. rapa*, where JA/ET pathway genes were enriched and SA-associated markers were largely uninduced at 7 dpi (Chu *et al*., 2014). Notably, classical TNL signaling genes (i.e., *EDS1*, *PAD4*, *ADR1*, and *ICS1*, etc.) were not activated in DH12075+*Rcr1* and in several cases were downregulated, further reinforcing that Rcr1 functions independently of canonical SA-mediated signaling during early infection. This supports a broader principle that TNL signaling is context-dependent, with root tissues favoring noncanonical outputs that bypass classical nodes. These findings align with recent studies proposing that *EDS1* and *NDR1* are not universally required for TNL function and may have tissue- or context-specific roles, particularly in roots (Mengiste and Liao, 2025).

Mechanistically, our data position Rcr1 as a noncanonical TNL that organizes immune responses through physically and functionally distinct modules (Fig. 3). Based on both interaction evidence and genetic validation, we define CP1, CP2, and AP as direct interactors of Rcr1 that are required for full resistance in *Rcr1*(+) backgrounds. These interactors modulate the expression of six downstream transcriptional responders, i.e., *WRKY33*, *WRKY55*, *ERF1*, *ERF109*, *EDS1*, and *NDR1*, whose transcript levels are altered in an *Rcr1*-dependent manner but without direct physical association. This dual-module structure, comprising a CP1–WRKY branch for recognition and an AP–ERF branch for signaling, forms the core of a modular defense system. CP2 contributes a minor supporting role, while canonical signaling components appear largely dispensable. Importantly, temporal dissection revealed that *Rcr1* activity is essential between 0 and 14 dpi, with the most critical period around 7 dpi. Together with the observation that *AP* knockouts attenuate *ERF1* and *ERF109* expression, this highlights that JA/ET signaling, particularly the ET arm, must be engaged early in the infection process. This timing aligns with broader evidence that immune pathway contributions shift dynamically across infection stages in clubroot (Liu et al., 2020; Wang *et al*., 2022). This modular architecture contrasts with paired NLR systems such as RGA4/RGA5 in rice (Cesari et al., 2013) or the NRC sensor-helper networks in Solanaceae (Wu et al., 2017), where downstream modules are also NLR-based. In the Rcr1 system, all downstream partners are non-NLRs, expanding the known signaling repertoire of TNLs and suggesting that combinatorial assembly of recognition and signaling elements could enhance immune plasticity (Cesari, 2018; Lee and Chae, 2020; Sun *et al*., 2020). The Rcr1 model thus provides a rare experimental example of modular NLR function in a natural crop-pathogen system and illustrates a conceptual framework in which immune recognition and signaling are separable but coordinated, enabling flexible outputs beyond canonical pathways.

From an applied perspective, our study highlights the importance of understanding not only effector specificity but also the architecture of resistance genes. The modular structure of Rcr1, with separable and functionally testable components, offers a blueprint for precision breeding strategies that stack orthogonal resistance modules. Such designs could improve durability by reducing redundancy and limiting the evolutionary escape routes available to the pathogen.

Rational deployment of modular NLRs, informed by mechanistic dissection, has the potential to create more resilient crop immune systems tailored to specific disease contexts. More broadly, the modular immune hub concept may help reframe how resistance genes are engineered, shifting emphasis from single-gene introgression toward rational assembly of complementary modules.

From a technical standpoint, this study leveraged two major innovations that enabled high-resolution mechanistic dissection. First, our NGS-Y2H approach enabled parallel screening of both TAN and LRR domains of Rcr1, allowing domain-specific interactome resolution. Compared to classical Y2H methods, which are limited by throughput and sensitivity, our pipeline prioritized high-confidence interactors based on enrichment scores and immune relevance. Although yeast-based systems underperform in detecting membrane-associated or transient interactions, our use of dual-domain baits increased specificity and coverage. Future studies will benefit from complementary methods such as co-immunoprecipitation or proximity labeling to capture more complete interactomes. Second, we developed a heat-inducible CRISPR/Cas9 system for *Rcr1* excision, providing temporal control over gene function during pathogen challenge. This system complements our dual-sgRNA mutagenesis approach used to knock out *CP1*, *CP2*, and *AP*, together forming a versatile toolkit for time-resolved and targeted gene manipulation in *B. napus*. This inducible platform also holds promise for future adaptation to tissue- or stage-specific editing, enabling precise functional analysis of immune components.

Despite these advances, several limitations must be acknowledged. First, this study focused exclusively on *P. brassicae* pathotype 3H, a dominant race in Western Canada. While this enabled clear mechanistic dissection, it remains unknown whether the same immune modules operate against other pathotypes such as 5X. Future comparative studies will be necessary to determine whether the Rcr1 configuration observed here is generalizable or pathotype-specific. Second, despite strong genetic and transcriptional support for the roles of CP1 and AP, biochemical validation of their effector interactions remains challenging, largely due to the fastidious biotrophic nature of the pathogen. Developing co-immunoprecipitation or *in vitro* binding strategies will be crucial not only to confirm these interactions at the protein level but also to clarify how *Rcr1* directly monitors pathogen effectors *in planta*. Finally, the conclusion that *Rcr1* bypasses canonical *EDS1*- and *NDR1*-dependent pathways is currently supported by gene expression evidence. Although the possibility of post-translational regulation of these components cannot be excluded, their consistently stable or reduced transcript levels, together with the lack of induction of other SA-pathway marker genes in resistant tissues, reinforce the model of a noncanonical signaling route.

In conclusion, this study identifies Rcr1 as the hub of a novel modular immune mechanism, providing an experimental demonstration that TNL receptors can achieve robust resistance through noncanonical, modular wiring. By coupling mechanistic insight with technical innovation, our work provides both a conceptual advance in plant immunity and a platform for rational resistance engineering and pathotype-resilient breeding strategies in canola.

## Data availability

All data supporting the findings of this study are publicly available. Raw NGS-Y2H data are deposited in the NCBI Sequence Read Archive (SRA) under BioProject ID PRJNA1306351. Two processed NGS-Y2H data tables, primer sequences, and one processed RNA-seq data table are provided in Supplementary Tables S3–S6 and published online with the paper.

## Funding

This research was supported by the Genomics Initiative of Agriculture and Agri-Food Canada.

## Author contributions

F.Y. and H.H. conceived the study and designed the experiments, H.H. conducted the experiments and performed the data analysis, H.H. and F.Y. wrote the paper.

## Declaration of interests

The authors declare no competing interests.

## Declaration of generative AI and AI-assisted technologies in the manuscript preparation

During the preparation of this work, the authors used ChatGPT in order to improve language. After using this tool, the authors reviewed and edited the content as needed and take full responsibility for the content of the publication.

## Supplementary Figures

**Fig. S1.**
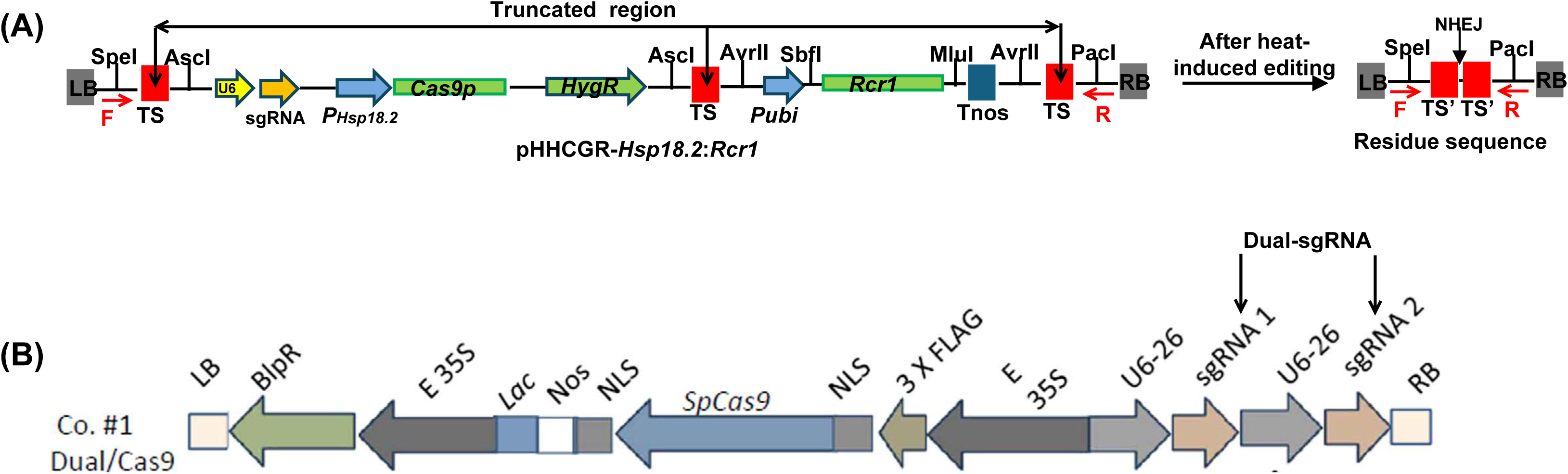
Vector construction. (A) Inducible *Rcr1* excision with heat treatment using pHHCGR-*Hsp18.2*:*Rcr1* (B) Knock-out mutation via Construct #1 with dual-sgRNA design (cited from Zhao et al., 2016)

**Figure S2.**
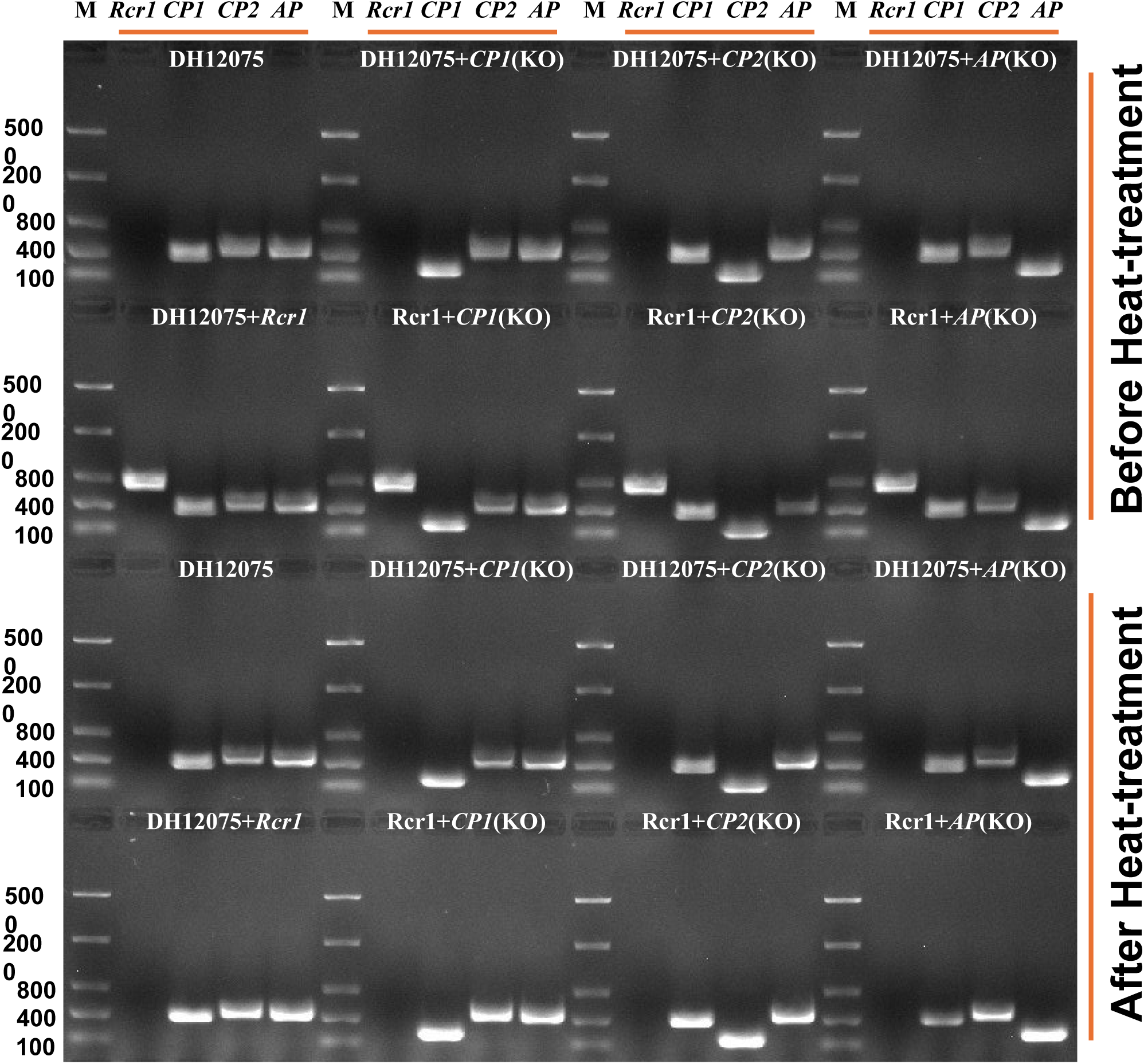
Plant genotype confirmation via PCR.

**Figure S3.**
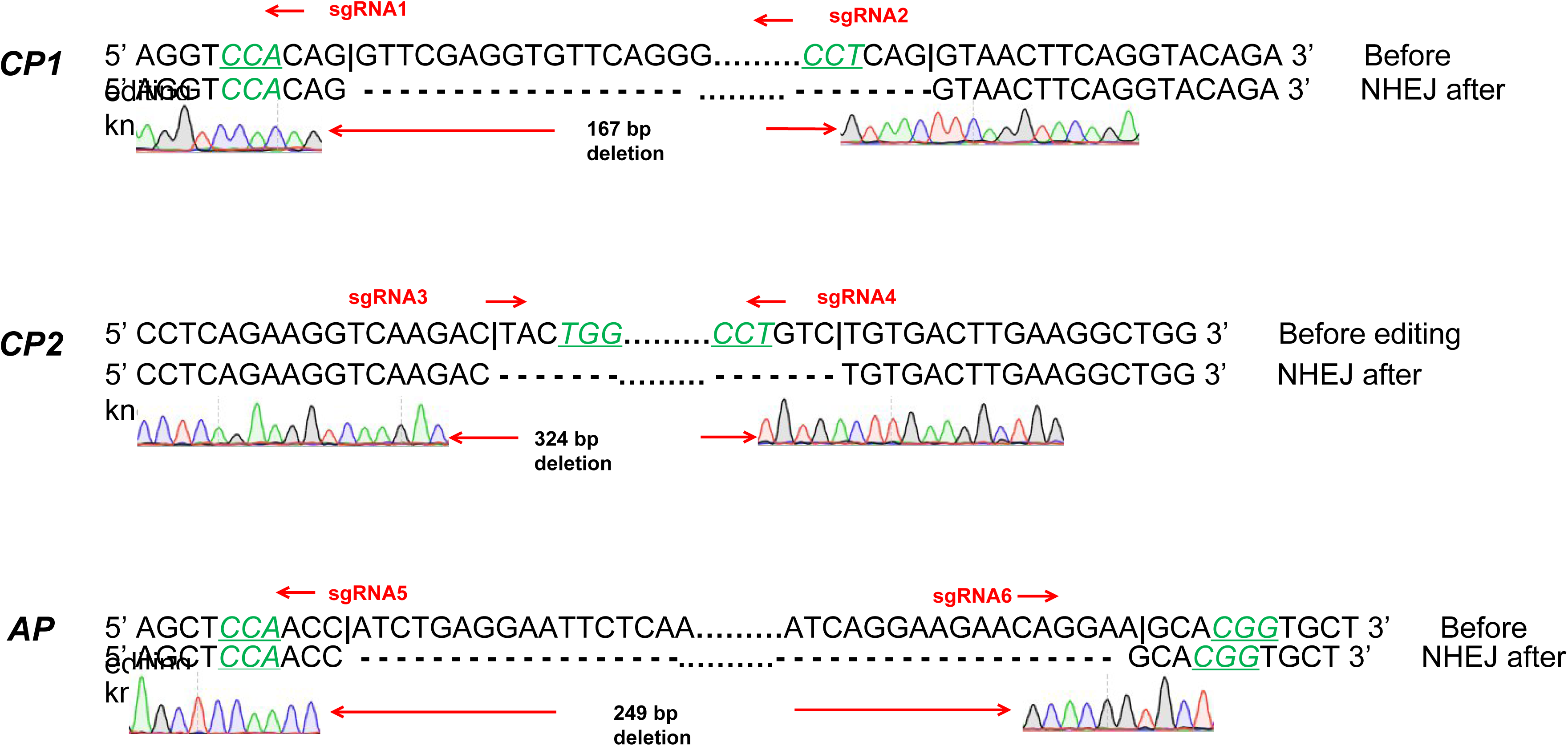
Plant genotype confirmation via Sanger sequencing.

**Figure S4.**
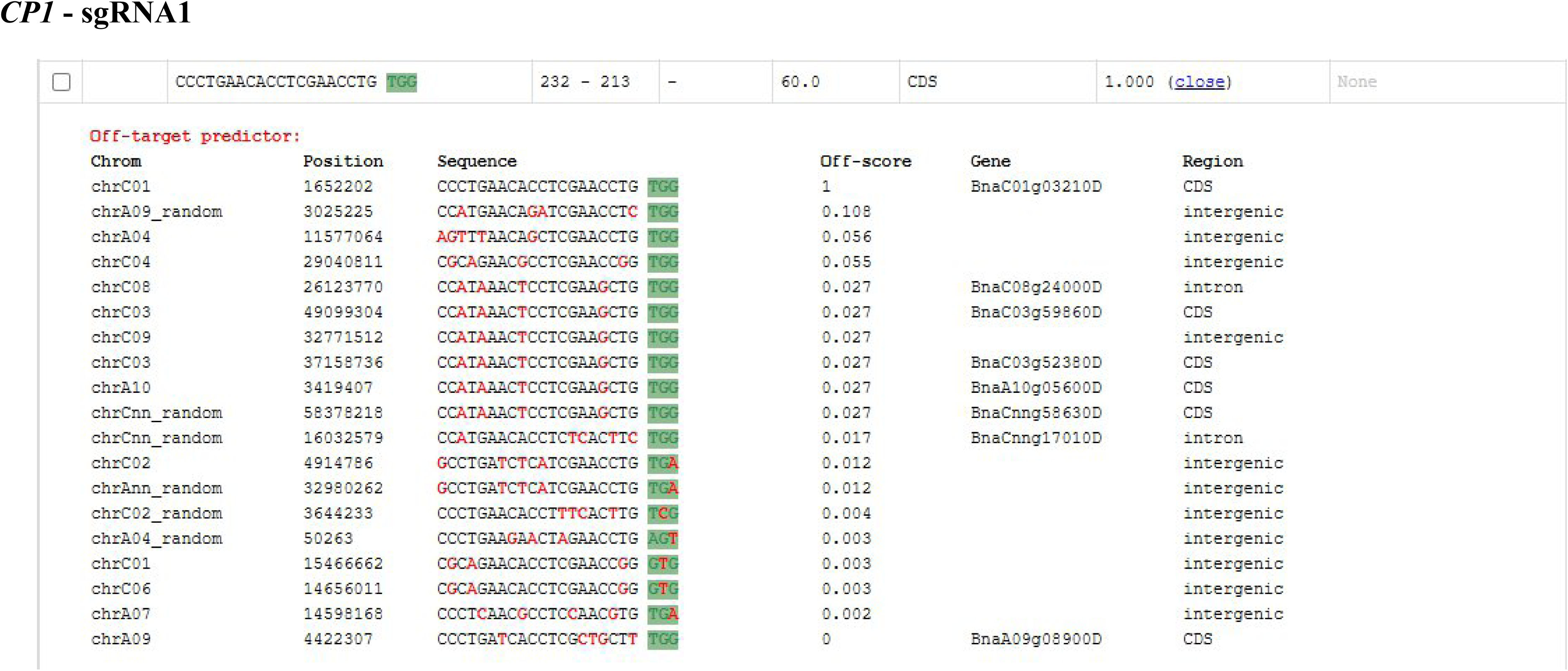
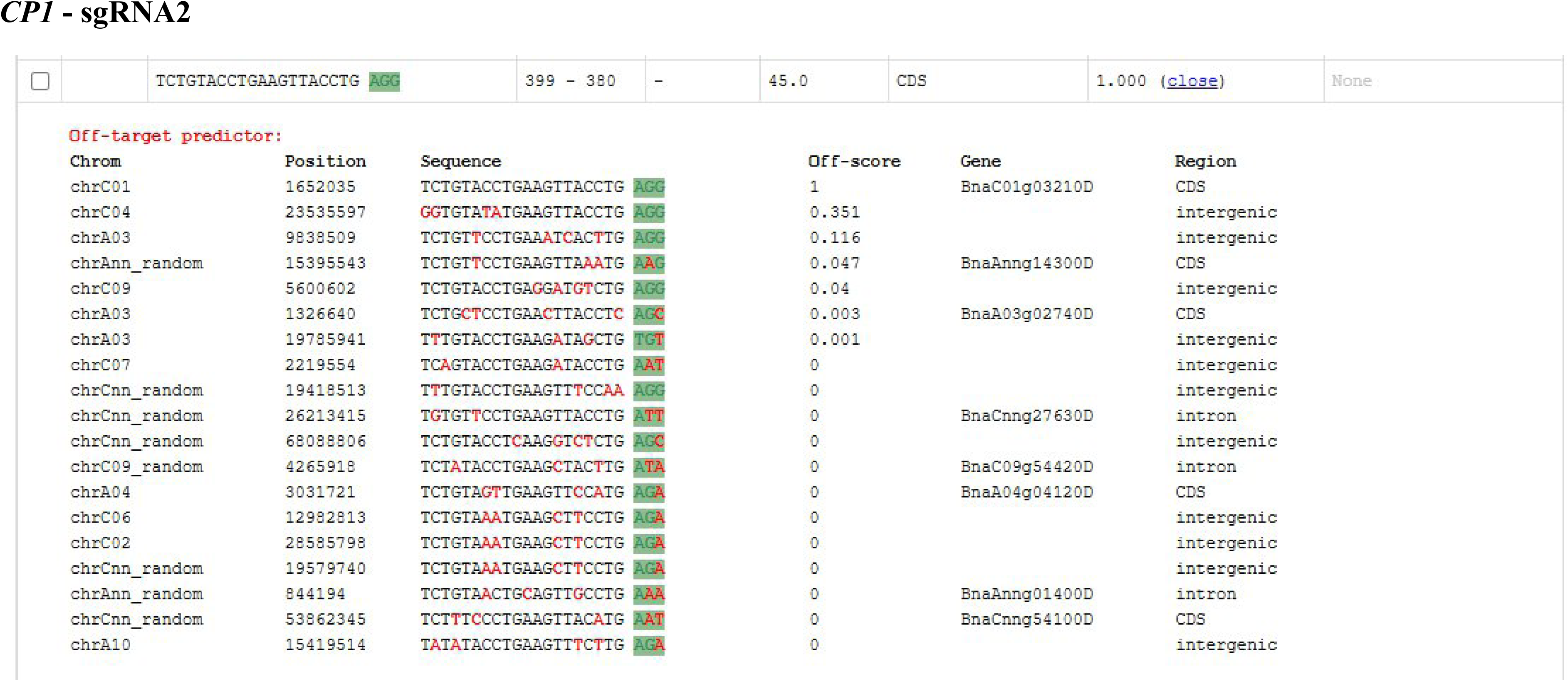
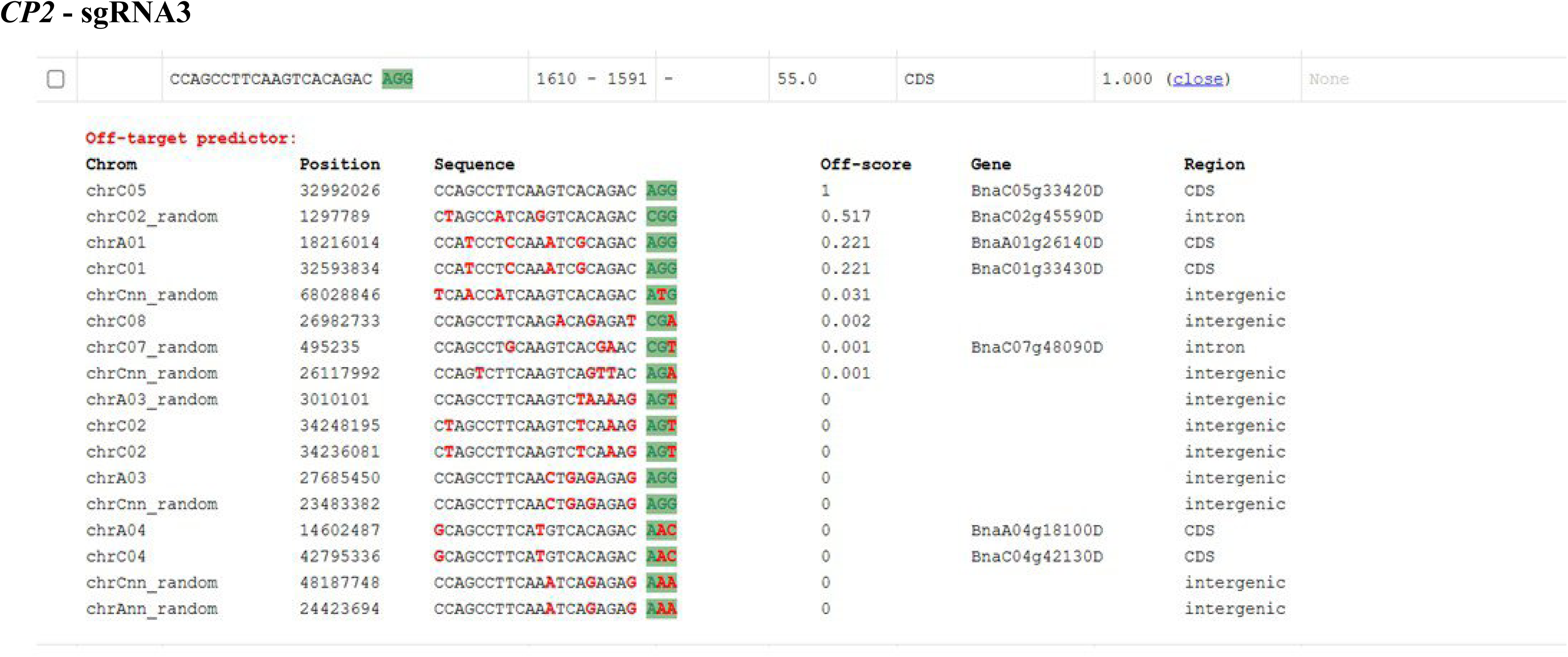
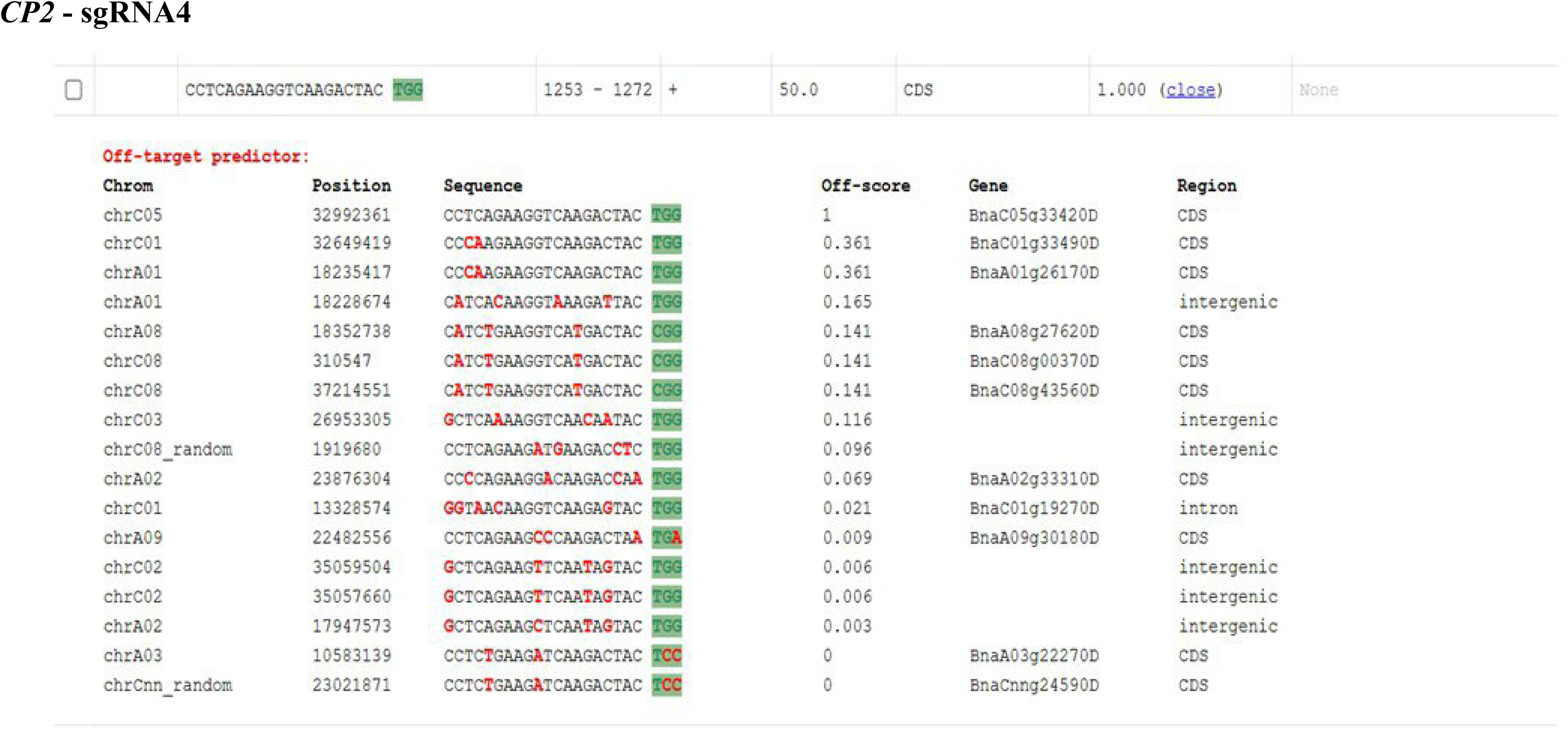
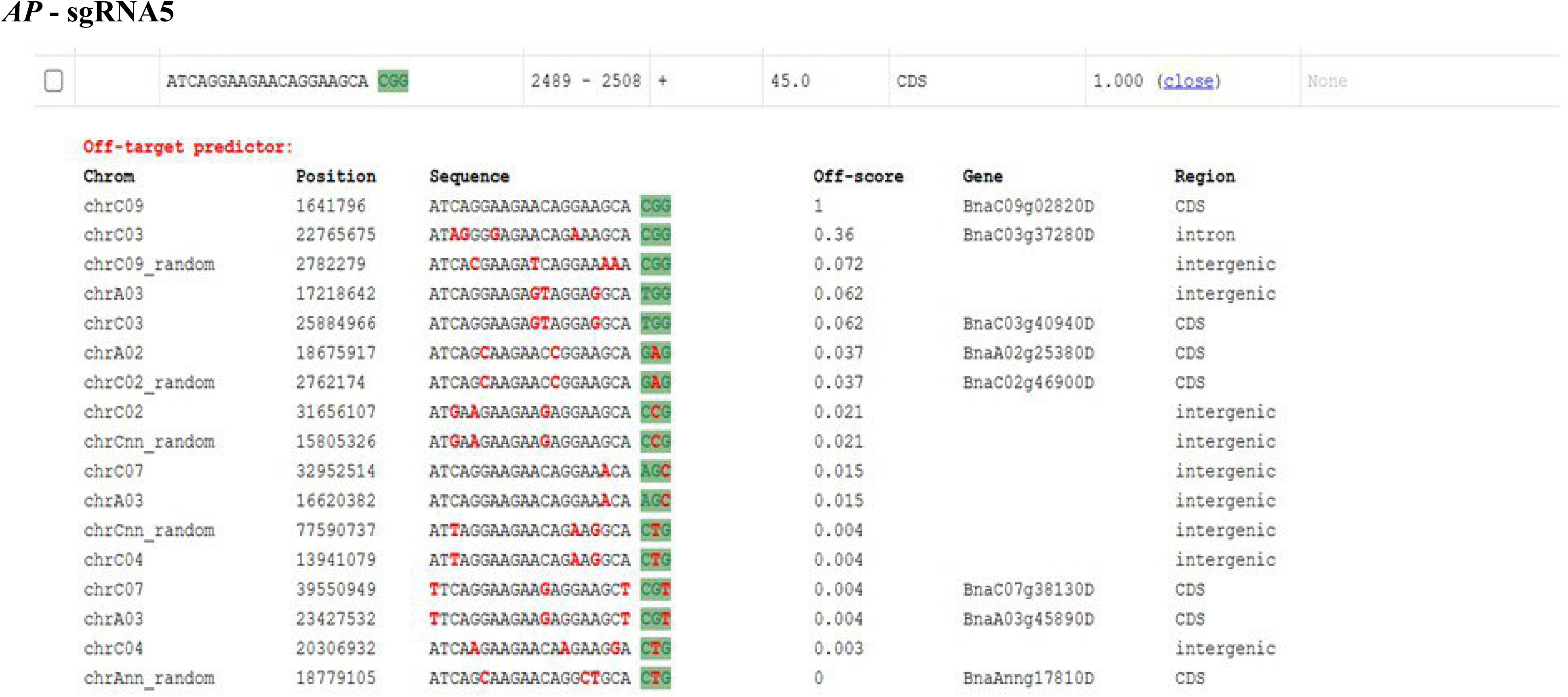
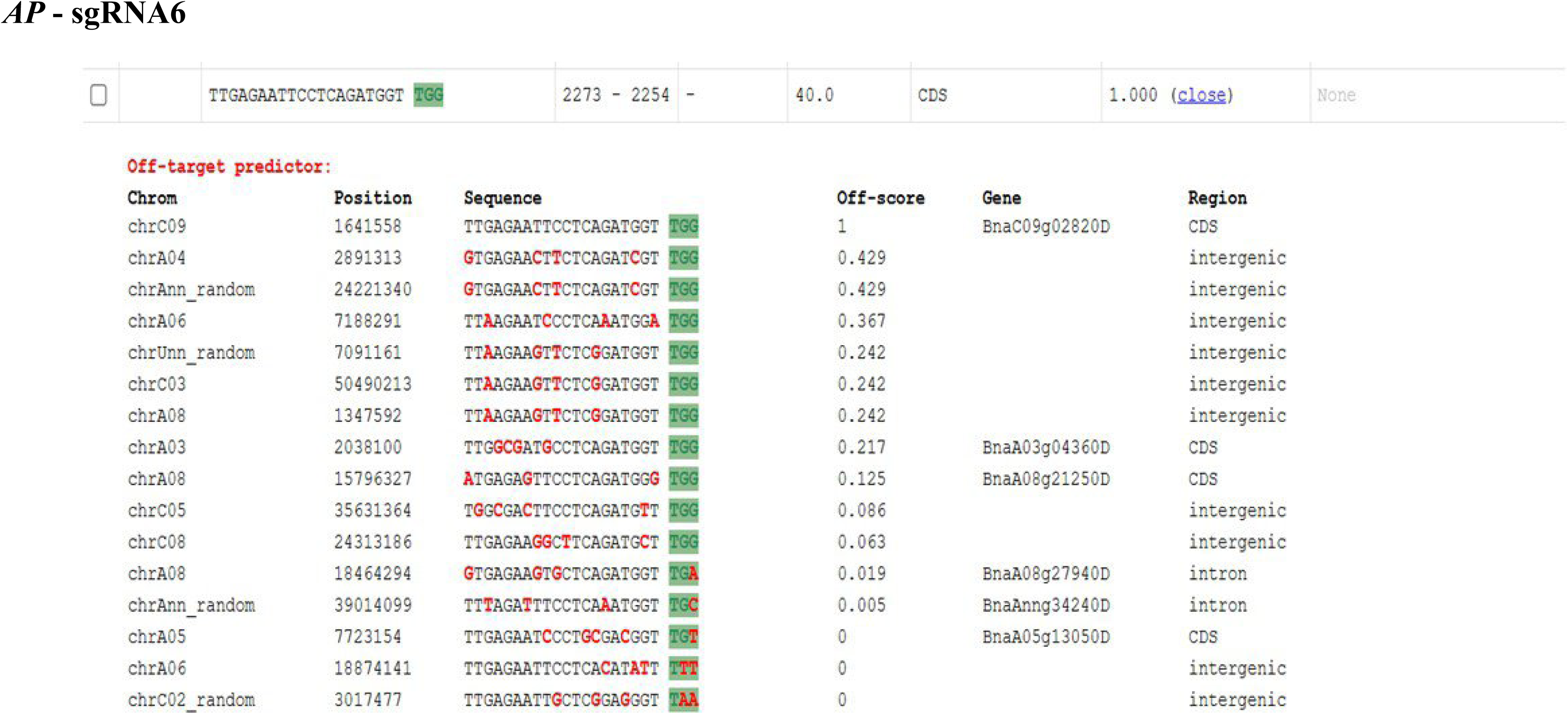
Off-target prediction from CRISPR-GE.

**Figure S5.**
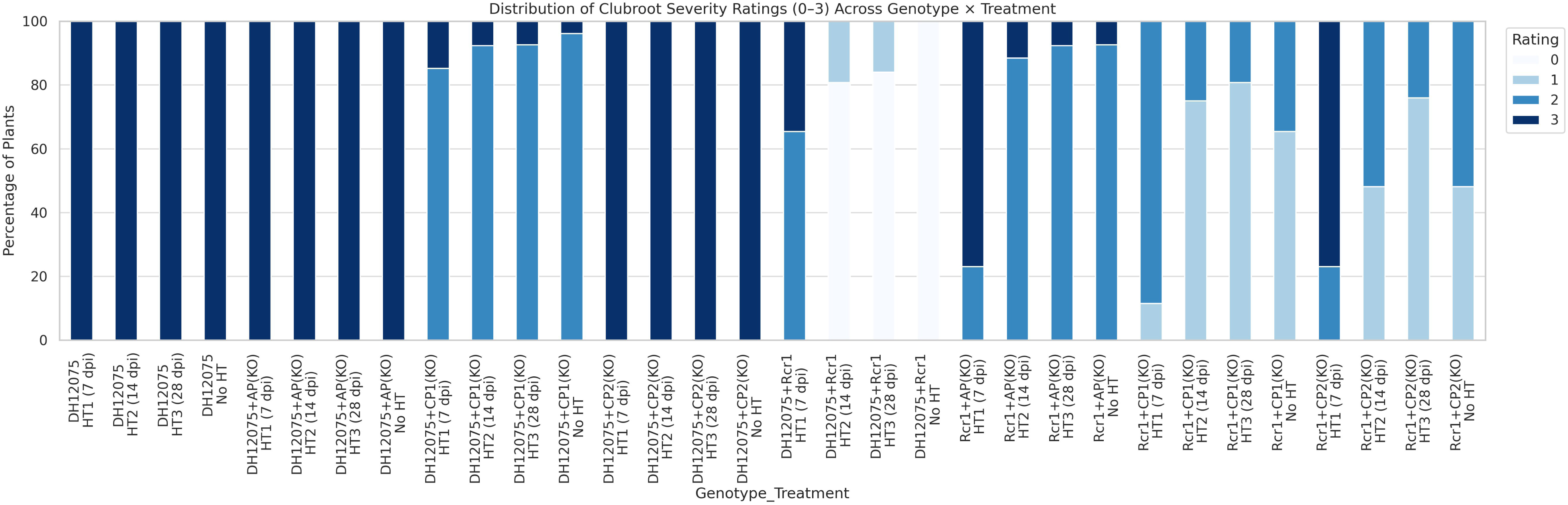
Distribution of clubroot severity ratings (0-3) across genotype x treatment.

**Figure S6.**
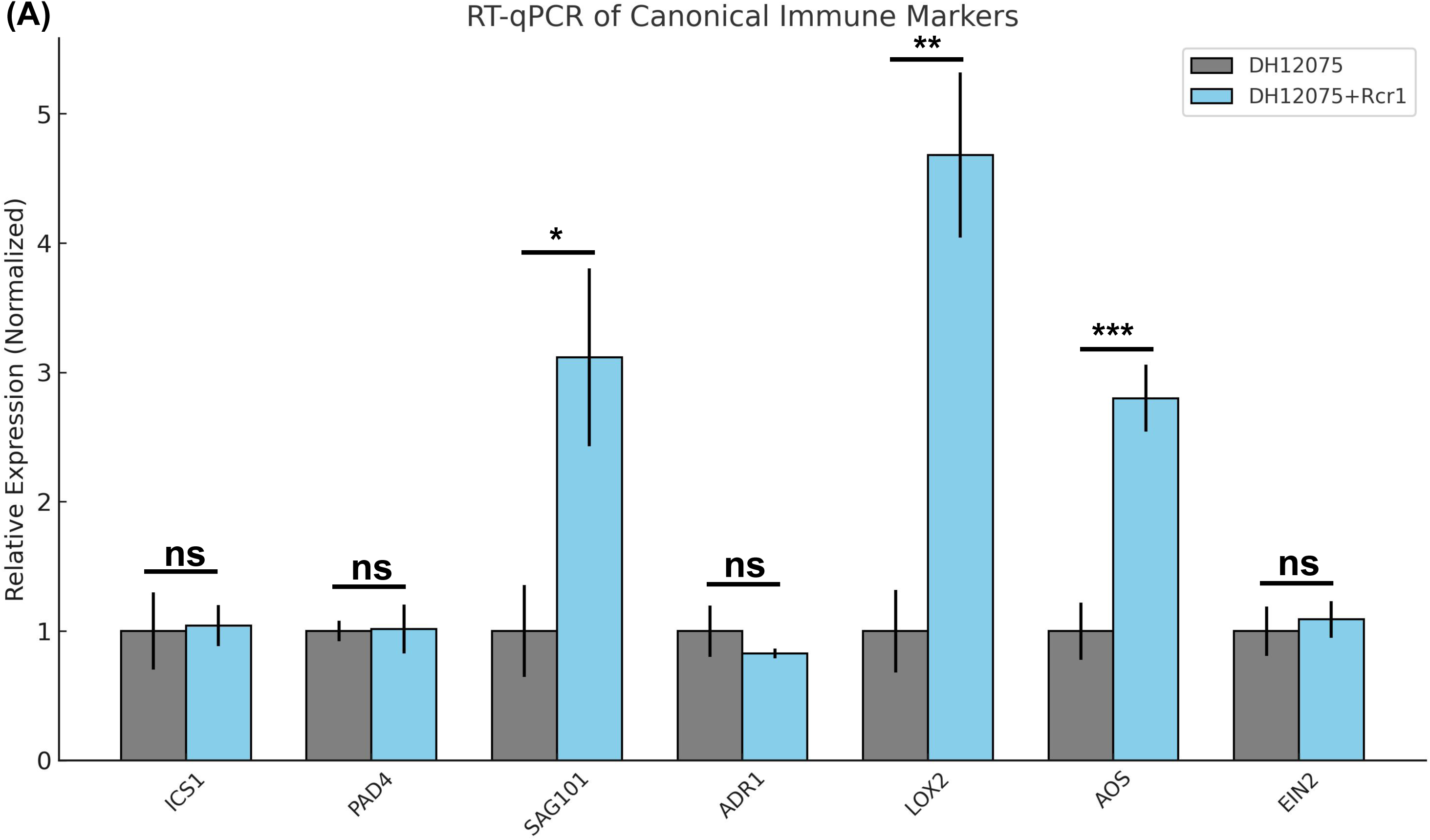
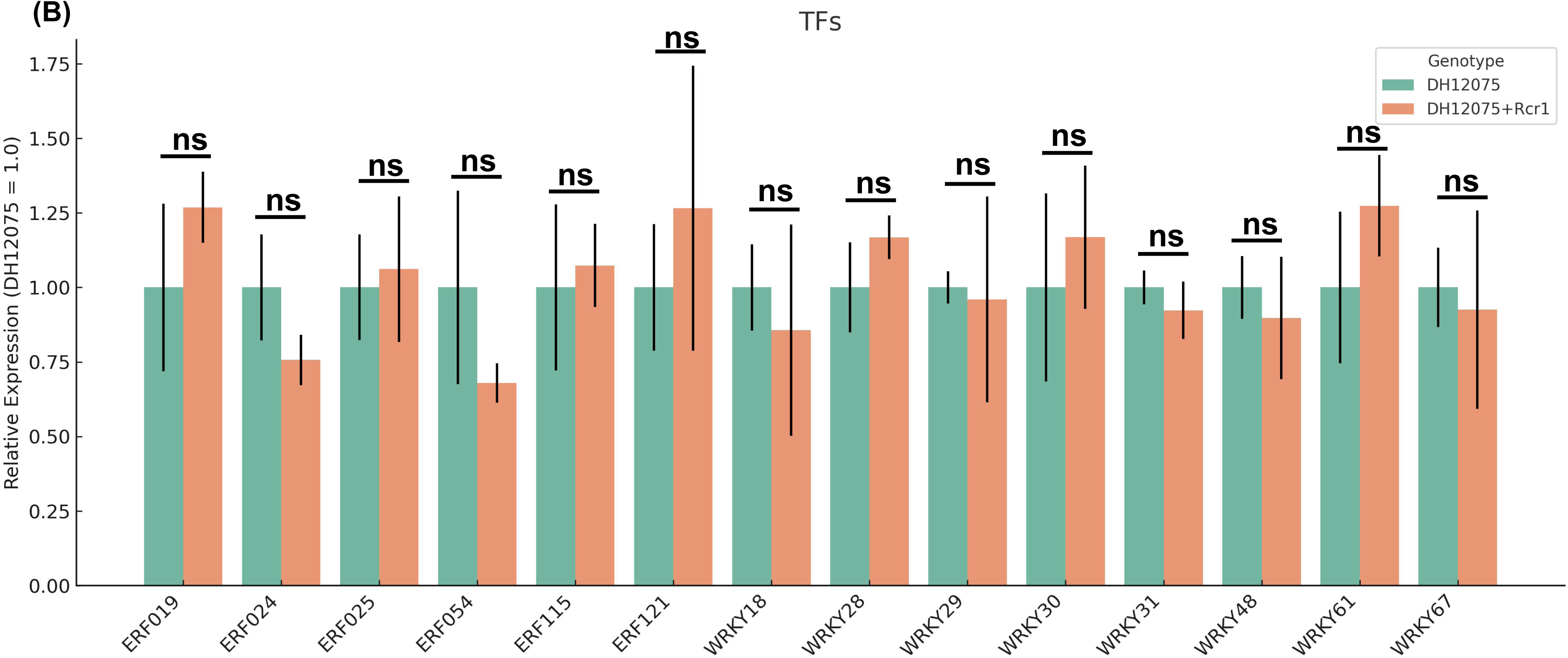
Gene expression profiling of *Rcr1*-related genes. (A) Canonical immune markers. (B)Transcription factors.

## Notes

http://www.ncbi.nlm.nih.gov/bioproject/1306351

### Competing Interest Statement

The authors have declared no competing interest.

